# De novo assembly and annotation of a highly contiguous reference genome of the fathead minnow (Pimephales promelas) reveals an AT-rich repetitive genome with compact gene structure

**DOI:** 10.1101/2021.02.24.432777

**Authors:** John Martinson, David C. Bencic, Gregory P. Toth, Mitchell S. Kostich, Robert W. Flick, Mary J. See, David Lattier, Adam D. Biales, Weichun Huang

**Author notes:** To whom correspondence should be addressed., Correspondence may also be addressed to Adam D. Biales.

## Abstract

The Fathead Minnow (FHM) is one of the most important and widely used model organisms in aquatic toxicology. The lack of a high-quality and well-annotated FHM reference genome, however, has severely hampered the efforts using modem ‘omics approaches with FHM for environmental toxicogenomics studies. We present here a *de novo* assembled and nearly complete reference of the fathead minnow genome. Compared to the current fragmented and sparsely annotated FHM genome assembly (FHM1), the new highly contiguous and well-annotated FHM reference genome (FHM2) represents a major improvement, having 95.1% of the complete BUSCOs (Benchmarking Universal Single-Copy Orthologs) and a scaffold N50 of 12.0 Mbps. The completeness of gene annotation for the FHM2 reference genome was demonstrated to be comparable to that of the zebrafish (ZF) GRCz11 reference genome. In addition, our comparative genomics analyses between FHM and ZF revealed highly conserved coding regions between two species while discovering much more compact gene structure in FHM than ZF. This study not only provides insights for assembling a highly repetitive AT-rich genome, but also delivers a critical genomic resource essential for toxicogenomics studies in environmental toxicology.

## INTRODUCTION

The fathead minnow (*Pimephales promelas*), found in most surface water across North America, can tolerate many harsh conditions. Its short lifespan (1-3 years) and high reproductive rate make it the most suitable species in fresh water for aquatic toxicology research and regulatory testing [1] among the most commonly used model species for ecological toxicity testing [2–5]. Consequently, there is a huge compendium of toxicity studies largely focused on apical responses, such as growth, mortality and fecundity, to chemical exposure. Though these studies have been critical in characterizing the risk posed by chemicals, they provide little insight into the underlying mechanisms determining organismal response. The stressor response phenotype is ultimately determined through the coordinated interaction of ‘omes layers (genome, epigenome, transcriptome, metabolome and proteome) [6], thus incorporating ‘omics endpoints into toxicity studies may provide key information regarding chemical mode of action (MOA) and factors driving the response phenotype [7].

Of the ‘omics, transcriptomics is perhaps the most well-studied due to the availability and maturity of experimental platforms, such as microarrays. Transcriptomic data have been successful in identifying chemical MOA, the etiology of disease, and a prognostic biomarkers [8–10]. The development of a FHM microarray has allowed researchers to evaluate across a large proportion of the transcriptome changes in gene expression [11–14]. The lack of comparable technological platforms for the other ‘omes has limited the characterization of their roles as a determinate of organismal response. As a result, interpretation of transcriptomic data is largely done without consideration of the other omics.

Over the last 10 years, advancements in next generation sequencing (NGS) have allowed ready access to other nucleic acid based ‘omics (epigenetics, genomics) [15]. Further, it has augmented transcriptomic data to include alternative splice variants and isoforms [16]. Epigenetics and genotype play significant roles in determining transcriptional levels, the development of disease, tissue specific responses, and alternative splice variants [7, 17, 18]. As the number of studies that incorporate epigenetics, the analysis of alternative splice usage and genomic sequencing has increased, it has become increasingly clear that solely focusing on the transcriptome provides only a glimpse of the total systemic response [19]. The inclusion of endpoints across other ‘omics provides additional layers of information that can be used individually or collectively to address gaps in our current understanding of the toxicological and stress responses, as well as the development of prognostic and diagnostic tools.

The availability of the first FHM genome assembly (dubbed here as “FHM1”), assembled with Illumina short-read sequencing, made it possible to use FHM for NGS ‘omics-based studies [20, 21]. However, due to the technical limits faced when assembling a genome from short read data the FHM1 assembly is highly discontiguous and incomplete, complicating the meaningful association of DNA and RNA sequence with gene expression so as to be useful for most NGS applications (e.g., transcriptomics). As interpretation of NGS results is highly dependent on the quality of the reference genome [22], there is a clear need for a high-quality reference genome. We here present a new, highly contiguous, nearly complete, and comprehensively annotated FHM genome assembly, which has been used successfully in-house studies with NGS-based technologies. The assembly was accomplished using a hybrid of long and short-read sequencing, optical mapping and Hi-C technologies. The quality of the new FHM assembly is comparable to, in terms of the complete BUSCOs and RNA-seq read mapping rate, the latest ZF GRCz11 assembly, as the latter represents the best characterized fish genome available [23].

## MATERIAL AND METHODS

### Source of specimens

All fathead minnows were obtained from the culture at the USEPA laboratory in Cincinnati, Ohio. This culture was established in 1998 and has since been maintained without any additional fish from the outside. Because the genome sequencing effort unfolded over several years, a few different fish were used for the different analyses.

### DNA library construction and sequencing

Brain and tail muscle tissue was isolated from a single 10-month-old male FHM. High molecular weight DNA was isolated using MagAttract (Qiagen, Germantown, MD). Samples were digested overnight using proteinase K. DNA concentration was checked on a Nanodrop ND-1000 spectrophotometer (Nanodrop Technologies, Wilmington, DE). Integrity of DNA was confirmed using a Bioanalyzer DNA 12000 chip (Agilent Technologies, Santa Clara, CA).

Long-read DNA libraries were prepared and sequenced by the Interdisciplinary Center for Biotechnology Research at the University of Florida (UF ICBR) on a PacBio RS II system. Sequencing was done in two rounds. For the first round of sequencing, BluePippin (Sage biosciences) was used to select templates greater than 20-kb, and libraries were sequenced on 24 SMRTcells using P6-C4 chemistry. For the second round of sequencing, the size selection was done using SageELF (Sage biosciences), and libraries were sequenced on 48 SMRTcells.

Short read sequencing libraries were prepared and sequenced by the Research Technology Support Facility (RTSF) Genomics Core at Michigan State University. Libraries were prepared using the Illumina TruSeq Nano DNA library kit. Sequencing was done in a paired-end 250-base format on both lanes of a dual flow cell in Rapid Run mode on an Illumina HiSeq 2500 System.

### RNA library construction and sequencing

RNA was isolated from multiple tissues from one adult male and one adult female fish, from unfertilized eggs, and from several developmental stages ranging from 24-h post-fertilization embryos to 30-d post-hatch larvae. Total RNA was isolated using Tri-Reagent (Molecular Research Center, Inc.) following manufacturer’s protocol and quantified using a Nanodrop ND-1000 (Thermo Fisher Scientific, Wilmington, DE). RNA quality was checked using RNA 6000 Nano Assay on an Agilent 2100 Bioanalyzer (Agilent Technologies, Santa Clara, CA). Two RNA pools were made: an adult RNA pool made up of equal amounts of RNA from each adult tissue, and a juvenile pool made up of equal amounts from eggs, embryos and larvae.

RNA libraries were produced and sequenced by the Research Technology Support Facility (RTSF) Genomics Core at Michigan State University. Separate libraries were prepared from the two RNA pools using the TruSeq Stranded mRNA LT sample prep kit (Illumina, San Diego, CA). Libraries were run in paired-end 150-base format on both lanes of a dual flow cell in Rapid Run mode on an Illumina HiSeq 2500 System. Base calling was done with Illumina Real Time Analysis software v1.17.21.3 and output was converted to FASTQ format with Illumina Bcl2fastq v1.8.4.

### Non-coding RNA library construction and sequencing

For sequencing of non-coding RNA (ncRNA), RNA was isolated from unfertilized eggs from a single female, from four 4-dph fry, from 20 6-dph fry, and from both brain and liver tissue from one adult male and one adult female. Samples were homogenized in Tri-reagent and processed using the Direct-zol RNA Microprep kit (Zymo Research; Irvine, CA) following manufacturer’s protocol. RNA quality was checked using an Agilent Bioanalyzer and RNA 6000 Nano chip. For each sample type, 1000 ng of total RNA was used as input for the TruSeq Small RNA Library preparation kit (Illumina) following manufacturer’s protocol. Libraries were sent to RTSF where they were pooled, checked for quality, and sequenced. The pool was loaded onto one lane of an Illumina HiSeq 4000 flow cell. Sequencing was performed in a 1×50bp single read format using HiSeq 4000 SBS reagents. Additionally, RNA was isolated from ten, 6-d post hatch larvae that had been exposed for 48 h to 10 mg EE2/l and ten unexposed larvae, following the procedures above. Libraries were made from the two sets of samples, and equal amounts of each were combined into one sample that was split and sequenced on two lanes of a flow cell.

### Generation of Bionano optical mapping data

DNA was isolated from blood of a six-month-old male FHM using the Blood and Cell Culture DNA Isolation Kit (Bionano Genomics, San Diego, CA). Nucleated fish blood was embedded in agarose plugs for titration in volumes of 1 μL, 3 μL, 8 μL, 20 μL, and 50 μL. A proteinase K digestion was performed, followed by additional washes and agarose digestion. The agarose plugs were sent to the McDonnell Genome Institute (MGI) Washington University for processing and analysis.

From this point, the DNA was drop dialyzed and allowed to equilibrate at room temperature for two days. The DNA samples were assessed for quantity and quality using a Qubit dsDNA BR Assay kit and CHEF gel. A 750ng aliquot of DNA was labeled and stained following the ‘Bionano Prep Direct Label and Stain’(DLS) protocol. Once stained, the DNA was quantified using a Qubit dsDNA HS Assay kit and run on the Saphyr chip. During the Saphyr run, the DNA were processed through a series of micro and nanochannels to prevent the linear DNA from folding back on itself. Once the DNA is linearized into the nanochannels, the data were imaged and captured by the Saphyr instrument.

### HiC library preparation and sequencing

Liver tissue from a single adult male of 10 - 12 months old was used for chromosome conformation capture Hi-C sequencing. Two HiC libraries were prepared in a similar manner as described previously [24]. Briefly, for each library, chromatin was fixed in place with formaldehyde in the nucleus and then extracted. Fixed chromatin was digested with DpnII, the 5’ overhangs filled in with biotinylated nucleotides, and then free blunt ends were ligated. After ligation, crosslinks were reversed, and the DNA purified from protein. Purified DNA was treated to remove biotin that was not internal to ligated fragments. The DNA was then sheared to ~350 bp mean fragment size and sequencing libraries were generated using NEBNext Ultra enzymes and Illumina-compatible adapters. Biotin-containing fragments were isolated using streptavidin beads before PCR enrichment of each library. The libraries were sequenced on an Illumina HiSeqX platform. The number and length of read pairs produced for each library was: 208 million, 2×150bp for library 1; and 220 million, 2×150bp for library 2. Together, these Dovetail HiC library reads provided 14,719.10 x physical coverage of the genome (10-10,000 kb pairs). This HiC library preparation and sequencing service was provided by Dovetail Genomics (Scotts Valley, CA) at https://dovetailgenomics.com.

### Genome Assembly

#### Assembling of the primary sequence contigs

PacBio subreads were extracted with the SMRT Link (version 6) *software tool* from PacBio. The extracted subreads were then corrected, trimmed, and assembled by the CANU assembler (version 1.8) [25, 26] to generate the first draft diploid FHM genome assembly (see Suppl Table 1 for parameter settings). Pilon (version 1.23) [27] was then used to polish this draft assembly by correcting miscalled bases, fixing mis-assemblies and filling small gaps in the assembly. Two sets of Illumina paired-end reads were used together for this polishing: 1) two lanes of 2X101 reads generated by Dupont from NCBI SRA (accession# SRR1304883 and SRR1301972), and 2) two lanes of 2X250bps from EPA. For each Illumina data set, BWA-MEM (version 0.7.15) [28] and SAMTOOLS (version 1.3.1) [29] were employed to align and sort Illumina reads to the assembly with BWA’s default paired-end read alignment protocol. After polishing, Purge_haplotigs (version 1.0.4) [30] was used to purge haplotigs, haplotype copies of primary contigs, from this diploid assembly. For a diploid genome, CANU often assembles both haplotypes as primary contigs for genome regions with a high degree of heterozygosity. This makes it necessary to purge such as haplotigs from this diploid assembly of FHM. This haplotig-purged assembly was then further polished with the same Illumina short reads by Pilon to correct misassembled contigs and fill gaps, some of which could be introduced by the haplotigs purging process.

#### Scaffolding of the primary sequence contigs

To generate chromosome-level genome assembly, we used the following three types of data to scaffold the primary sequence contigs assembled from PacBio reads: 1) two sets of long Illumina mate-pair or jump-library sequencing reads, which were downloaded from the public NCBI SRA database including one 40Kb fosmid library (SRR1304690) and one 6Kb mate-pair library (SRR1301983), 2) HiC Illumina paired-end sequencing reads (see *HiC library preparation and sequencing* section for the details), 3) optical mapping data generated using Bionano Direct Label and Stain (DLS) technology (see *Generation of Bionano optical mapping data* section for the details).

To achieve the best scaffolding results, we scaffolded the assembly of primary contigs four times in the following order: 1) scaffolding using the HiC library data, 2) scaffolding using the Illumina mate-pair library data, 3) hybrid scaffolding with Bionano optical mapping data, and 4) final scaffolding with the HiC library data. For both assembly scaffolding with HiC data, SALSA (version 2.2) [31] was used as the scaffolding tool and BWA-MEM (version 0.7.15) was the alignment tool. For the first HiC scaffolding, reads were mapped to the assembly with the default BWA-MEM paired-end read mapping protocol while reads were aligned with the option “-SPM” to skip mate rescue, read paring, and mark shorter split hits as secondary for the final HiC scaffolding. SSPACE (Standard version 3.0) [32] tool was used for the second assembly scaffolding with the Illmina mate-pair library data. SSPACE was run with the option “-x 0 -m 32 -o 20 -k 5 -a 0.70 -n 15 -p 1 -v 0 -z 0 -g 0 -S 0”. The hybridScaffold.pl tool in the Bionano Solve (version 3.3, https://BioNanogenomics.com/support/software-downloads) was employed to carry out the hybrid scaffolding with Bionano optical mapping data. It was run with the option “-B 2 -N 1” to ensure no cut to the sequence genome assembly while allowing cuts to Bionano genome map when there are conflicts between two genome assemblies.

For both genome assembly and scaffolding, we tested several different tools and tried with different parameter settings to find the best tools and running options to achieve the best FHM assembly. For example, we tested CANU (multiple versions), FALCON (version 1.2.4) and FALCON-UNZIP (version 1.1.4) in the pb-assembly pipeline (version 0.0.6) [33], DBG2OLC [34], MARVEL assembler [35], and Minimap/Miniasm [36] as well some other assemblers. We also tested different running assembly options, particularly for CANU, for both diploid and haploid assemblies. For assembly scaffolding, we tested several scaffolding tools, e.g., SSPACE, BESST [37], BOSS [38], OPERA_LG [39] for scaffolding with Illumina mate-pair reads. In addition, we tried different alignment options and different scaffolding orders with different types of data for scaffolding. For example, we tried scaffolding first with the Illumina mate-pair data, then Bionano, and finally HiC data, but it turned out to be not the best.

### Genome assembly evaluation

We evaluated a genome assembly in multiple ways. We first checked if genome assembly size was closed to the estimated size from other sources e.g., flow cytometry, and compared N50 for assembly continuity of different genome assemblies. We also used mapping rates of short Illumina reads from genome resequencing and RNA-seq generated in house to assess completeness of the genome assembly. Finally, we used BUSCO (version 3.0.2) [40, 41] to evaluate genome completeness and duplication level of individual assembles.

### Identification of transposable and repeat elements

The transposable elements (TE) in the FHM genome were identified with a combination of de novo finding and homolog-based search approaches. For the *de novo* TE finding, we used RepeatModeler (version 2.0.1) [42] and RepeatMasker (version 4.1.0) (http://www.repeatmasker.org) to construct a *de novo* FHM-specific repeat consensus library with TE classification. In this step, RepeatScout (version 1.0.6) [43] and RECON (version 1.08) [44] were deployed to discover all TE elements and generate TE family models. Additionally, RepeatModeler LTR module was used to carry out an independent run to identify and classify long terminal repeats (LTR). Specifically, the LTR module was run with LTRharvest from GenomeTools (version 1.5.9) [45] for LTR discovery and annotation, LTR_retriever (version 2.8) [46] for LTR filtering and annotating, and CD-HIT (version 4.8.1) [47], MAFFT (version 7.453) [48], and Ninja (0.95-cluster_only) [49] for LTR clustering and family classification. All discovered TE families were then consolidated into a single FHM-specific TE library by clustering and removing redundancy by the RepeatClassifier module of RepeatModeler.

Potentially non-TE protein coding sequences in this FHM-specific TE library constructed by RepeatModeler was then filtered out by the following procedure. First, we obtained the known protein sequences of the FHM genome from the UniProt Knowledgebase (UniProtKB), which yielded only 283 protein sequences. To expand the protein dataset, we combined it with the uniprot zebrafish genome protein set (62,372 sequences) for a total of 62,655 sequences. We then removed known TE found in the RepeatMasker TE database from the protein sequence set by searching with BLASTP (NCBI-BLAST version 2.10.0+) and filtered out all hits with an E-value <1e-5. After that, we searched potential non-TE coding sequences in the FHM-specific TE library by BLASTing against this TE-filtered protein dataset and filtered out non-TE sequences from this FHM-specific TE library of hits with an E-value < 1e-5.

Using this FHM-specific TE library together with the existing repeat databases including the Dfam and RepBase, RepeatMasker (version 4.1) (http://www.repeatmasker.org) was used to carry out the whole-genome repeat analysis of this FHM genome assembly, identifying all repetitive regions including low-complexity regions and simple repeats. Specifically, RepeatMasker was run under the sensitive mode (-s) with the searching engineer RMBlast (version 2.10.0+) from http://www.repeatmasker.org, and Tandem Repeats Finder (version 4.09) [50]. The two known repeat databases used for the analysis were: 1) Dfam (version 3.1 June 2019, 6915 entries), an open database of transposable element (TE) profile HMM models and consensus sequences [51], and 2) RepBase (RepeatMaskerEdition-20181026) from Genetic Information Research Institute (http://www.girinst.org), which is an interspersed repeat database including all representative repetitive sequences from eukaryotic species [52].

### Gene prediction

#### Ab initio prediction

The ab initio gene prediction was carried out by the latest version of AUGUSTUS (Version 3.3.3) [53], which produces an ab initio gene prediction based on a Generalized Hidden Markov Model (GHMM). For the ab initio gene prediction, RepeatMasker was used to generate soft-masked version of the FHM genome assembly, and AUGUSTUS was then run for prediction using the built-in zebrafish model on both DNA strands (parameter setting --strand=both --softmasking=1 --species=zebrafish --genemodel=partial).

### Gene prediction with RNA-Seq data

RNA-seq reads were used to help define coding gene regions and exon boundaries. In order to facilitate this process, we assembled the RNA reads using Trinity [54] in genome-guided mode and on *de novo* mode. First, sequence reads were error corrected using rCorrector (version 1.0.3.1) [55]. Next, read pairs with one or both reads flagged as ‘unfixable_error’ by using FilterUncorrectabledPEfastq.py (github.com/harvardinformatics/TranscriptomeAssemblyTools) were removed. TrimGalore (version 0.6.1) [56] and the Cutadapt [57] were then used to remove adapter sequences from the RNA reads at the cutoff of stringency 1 and evalue 0.1. Trinity (version 2.5.0) was employed for both *de novo* and genome-guided assembly of the RNA-seq reads. For the genome-guided assembly, STAR (version 2.6.0a) [58] was used in two-pass mode to map processed RNA-seq reads to the genome assembly. Two annotation pipelines, Maker [59] and PASA/EVidenceModeler (EVM) [60] were employed to predict protein coding gene models.

#### Annotation pipelines

Two iterations of Maker (version 2.31.9) were run first, using as inputs the genome sequence, a repeat library identified with RepeatModeler (version 1.0.11), the Trinity genome-guided RNA assembly, ~22.5K non-redundant FHM ESTs and a set of reference proteins. The FHM ESTs were obtained from the Joint Genome Institute (JGI). The reference proteins were downloaded from the European Bioinformatics Institute (EBI) and they were from the following 17 species: *A thaliana, B subtilis, B floridae, C elegans, C intestinalis, D rerio, D discoideum, D melanogaster, E coli, G gallus, H sapiens, M musculus, L oculatus, O latipes, S cerevisiae, S pombe, and X tropicalis*. Augustus [61], SNAP [62], and GeneMark-ES [63] were all used for gene prediction, employing FHM-specific hidden markov models (HMM) trained with gene models obtained from a preliminary Maker run. SNAP was retrained for the second iteration using all the gene models output from the first iteration of Maker. PASA (version 2.3.3, employing blat (version 36.1) [64] was used together with gmap (version 2017-11-15) [65], TransDecoder (version 5.5.0) [66], and EVidenceModeler (EVM, version 1.1.1) as an alternative path to produce protein coding gene models. Inputs to EVM included gene predictions and alignment information generated during the Maker runs, and gene/transcript predictions and alignments from PASA. Maker and TransDecoder models producing proteins with perfect BLAST alignments to reference proteins were included as highly weighted inputs to EVM. EVM output models were processed back through PASA (two iterations) in annotCompare mode to add UTR sequence and/or generate alternative transcripts. The software tools employed as part of Maker and PASA/EVM are listed in the Suppl Table 2.

#### Selection of preferred model set and additional processing through Maker

Transcript and protein models from both pipelines were assessed using BUSCO [40]. The 37,190 PASA/EVM models exhibited ~11% more complete BUSCOs than the Maker models and were selected as the preferred model set (Suppl Figure 1). Mapping rates for the two model sets were also compared using ~6.43 (about 2.5%) million randomly selected paired-end RNA reads by USEARCH (version 9.2) [67] from the full RNA-seq dataset of ~320M pairs of 150bp paired-end reads, which had been adapter/quality trimmed by BBDuk (version 37.41) [68]. The sampled reads were mapped by STAR to transcripts corresponding to the longest model protein sequences (exemplars) from any given gene. Mapping rates were equivalent even though there were ~17% fewer models in the Maker set, so we elected to run the PASA/EVM models back through Maker as input predictions attempting to add additional UTR information to boost per gene mapping rates. Due to not invoking the “keep_preds” parameter during the Maker run only ~69% of the input PASA/EVM models were returned, so original PASA/EVM models exhibiting no coding sequence overlap with the new Maker models were returned to the model set. The unique mapping rate increased by ~6% in the modified set relative to the input set, while BUSCO scores remained the same.

#### Gene model filtering

36,881 models remained, which was higher than the number of genes expected based on the number of genes in zebrafish. To address this, models were filtered using a process based on mapping evidence and homology to reference proteins. Transcripts representing complete single-copy BUSCOs were identified, and Bowtie2 (version 2.3.5.1) [69] was used to map, in unpaired mode, the full dataset of RNA-seq paired-end reads to exemplar transcript models. The tenth percentile of correct strand mappings/total mapping was calculated for the exemplar transcripts representing single-copy BUSCOs, and the resulting value was 0.842. This value was used as a cut-off for a 2-sided lower 95% CI binomial estimate of the ratio of correct strand counts/total counts for non-BUSCO genes (transcripts); genes with an estimated lower CI boundary >=0.842 were retained in the final models. A homology criterion was also applied to evaluate models that may be valid but that represented genes that were not highly expressed in our data. Diamond (version 0.9.22.123) [70] was used to do a BLASTX search where the query sequences were exemplar transcript models meeting the mapping criterion above but that were not identified as single-copy BUSCOs, and the targets were our EBI reference proteins. Using only positive frame alignments, the tenth percentile of percent ID to the reference proteins was 0.503, and the tenth percentile of percent overlap to the best hit reference protein was 0.404. These values were used as homology cut-offs. Models that met both homology criteria were also kept as the final models.

#### Manual curation of select models

The original EVM protein sequences were compared to their corresponding proteins in the 26,150 filtered gene models (26,150 genes resulting in 47, 578 proteins) to identify any that had been changed as a result of being run back through Maker. Any that had changed had both their original EVM protein sequence and post-Maker protein sequence BLASTed (version 2.6.0+) [71, 72] against our EBI reference proteins and the coverage to both query sequence and subject sequence of the best hits was assessed. Original EVM proteins exhibiting better coverage were restored to be part of the final protein sequences. To maintain mapping improvements (i.e., UTR additions) that might have been gained by the passage through Maker, the restoration was done by manually adjusting CDS and/or UTR start/stop coordinates and/or CDS phase as necessary in the corresponding Maker transcript model so that the CDS segments/phases mirrored those in the original EVM models; any additional sequence segments added by Maker were designated as UTR sequence. Based on this assessment, a relatively small number (303) of proteins were manually adjusted.

#### Gene annotation assessment

We benchmarked our gene prediction models for the FHM by comparing FHM2’s BUSCO scores and RNA-seq read mapping rates to ZF. For the analysis, we downloaded the ZF reference genome (GCF_000002035.6_GRCz11_genomic) and its gene annotation gff file from NCBI at https://ftp.ncbi.nlm.nih.gov/genomes/all/GCF/000/002/035/GCF_000002035.6_GRCz11. For the comparison to FHM, we extracted the 992 ZF scaffolds with ‘Primary Assembly’ in their definition lines (992) and the single sequence dubbed “mitochondrion, complete genome” from the ZF sequence file and also extracted the corresponding scaffold records from the ZF gff file. Transcript and protein sequences were then produced from the extracted gff file using gffread (v0.11.4) from https://github.com/gpertea/gffread from [73]. Transcripts without corresponding proteins were removed, so the final numbers of transcripts and proteins was the same, 47,834.

To perform the mapping rate comparison, ~326M paired-end ZF RNA-seq data were downloaded from NCBI’s short read archive (SRA) (see Suppl Table 3). The RNA-seq reads selected for download represented different tissues and life stages of ZF roughly mimicking the total number and tissue composition of our in-house FHM reads. All reads in the downloaded datasets were trimmed to 50 bases, the minimum read length in any of the sets; reads pairs where either read was <50 bases were discarded. From each trimmed dataset, five different random samplings, each representing 5% of the sequences in that dataset using USEARCH. The reads selected from each iteration of USEARCH from each downloaded dataset were then combined, resulting in the end in five datasets of randomly selected reads, each with ~16M 50 bp paired-end reads. These five sets of ~16M reads were them mapped, as both paired-end and single-end, with STAR (version 2.7.1), to the 62,884 extracted transcripts (47,834 protein-coding) from the ZF primary assembly. Default STAR parameters were used with the exception of “--outFilterMultimapNmax 20”. Mean mapping rates across the five datasets were calculated for both the paired-end and two single-end mapping runs. For comparison, we also trimmed our ~320M 150bp paired-end FHM RNA-seq reads to length 50 and again used USEARCH to randomly select five sets of 16M reads from the trimmed reads. The reads were then mapped to the 47,578 FHM2 transcripts in the same manner as the ZF reads had been mapped. As an additional comparison between the two species BUSCO (version 3.1.0) analyses were performed on both transcripts and proteins from FHM2 and ZF using BUSCO’s Actinopterygii reference set.

### Identification of short non-coding RNAs

FHM tRNAs were identified using tRNAscan-SE (v. 2.0.5), employing the program’s EukHighConfidenceFilter for a high confidence tRNA gene set (i.e., estimated most likely to be involved in ribosomal translation) [74]. Comparisons of high confidence tRNA gene sets across species used data from GtRNAdb2 (http://gtrnadb.ucsc.edu).

FHM microRNAs (miRNA) were identified using the command-line version of miRDeep*, MDS_command_line_V37 [75]. Small RNA-Seq reads for the miRDeep* run were pooled (1.07 billion reads total) from two separate FHM sources: 1) smRNA-Seq reads from FHM 48-hour post-hatch larva that had been statically exposed for 48 h to 0 (10 samples) or 10 ng ethinylestradiol/l (10 samples) (694 million reads sub-total), and 2) four separate samples of adult FHM liver (both sexes mixed), brain (both sexes mixed), unfertilized eggs from a pre-spawn female, and a pool of 4 96 h post-hatch fry (377 million reads sub-total). miRDeep*’s MD.jar was run under default conditions (Min score: -10; miR length: 18-23) and using the TruSeq (RA3) adapter sequence TGGAATTCTCGGGTGCCAAGG. The score is the log-odds probability of a sequence being a genuine miRNA precursor versus the probability that it is a background hairpin. The resulting “mature sequences” were sorted by score and duplicate sequences removed to yield 11,069 unique miRNAs. These miRNAs were further sorted by score and probed for exact matches with 347 *D. rerio* miRNAs from mirbase.org (release 22). The sorted miRNAs were also analyzed by BLASTing against the general mirbase.org cross-species database to evaluate change of E-value distribution at different score cutoffs. The final miRNA prediction results were based on the stringent cutoff 100, which was selected by testing different cutoffs on impact on the number of predicted miRNAs exactly matched to those reported in the ZF.

FHM piwi-interacting RNAs were identified using two separate programs: Piano [76] which predicts transposon-related piRNAs and piRNN [77] which identifies piRNAs globally by using a neural network piRNA prediction model. Piano uses a Drosophila-trained model for prediction and requires a species-specific transposon file as well as a smallRNA file. piRNN provides the code to develop a species-specific prediction model trained with species-specific piRNA (along with miRNA and tRNA for negative controls). piRNA sequences for this development came from a combination of FHM Piano output and ZF piRNA sequences to which FHM smRNA sequences aligned. Trimmomatic-trimmed smRNA sequences were then used for the primary piRNN python program to predict global FHM piRNN sequences. piRNN output was further processed to enable removal of sequences which aligned with FHM miRNAs and removal of sequences smaller than 25 nucleotides and greater than 40 nucleotides.

A general analysis of FHM short non-coding RNAs was done using Infernal-1.1.2 [78], employing the Rfam (v 14.0) library of covariance models, to search for homologues to known non-coding RNA families (e.g., small nuclear RNA (snRNA), small nucleolar RNA (snoRNA), tRNA, ribosomal RNA (rRNA), miRNA).

### GO terms and KEGG pathways annotation

Gene functional annotation was based on both sequence homology found BLAST search and protein domains and motifs identified by InterProScan [79]. The NCBI BLAST (version 2.10.1) [72] was used to search all predicted FHM protein sequences against the NCBI landmark proteome database, which consists of taxonomically diverse non-redundant set of proteins from 27 best annotated model organisms spanning a wide taxonomic range including zebrafish, human and mouse. We used the “*subject_besthit*” option for the BLASTP search and kept only hit sequences with <BLAST e-value 10^−5^. BLAST results were then entered into Blast2GO (version 5.2.5) (https://www.blast2go.com)[80], which retrieves GO terms associated the homologous BLAST hits. GO terms associated to only the top 20 homologous hits were kept. For GO assignment and annotation, we first used BLAST e-value 10^−6^ as the cutoff to filter out mapped GO terms, and then used the Blast2GO’s default annotation rule to assign GO terms to each query protein sequence. The cutoff of the Blast2GO annotation score, which is based on the sequence similarity score, weights of evidence codes, and number of children GO terms, was 55 for the GO term annotation.

Independently, we ran InterProScan (version 5.45-80) to retrieve domain/motif information in a sequence-wise manner for each predicted FHM protein sequence. We then loaded the InterPro annotation results into Blast2GO and obtained the GO terms associated with InterPro annotation. InterPro-based GO term annotations were then merged with the GO term annotations from the BLAST homolog-search using Blast2GO’s augmentation and annexing processes to generate the final full GO annotation data. An abbreviated version of the GO annotation data, GO slim, containing a subset of terms, which is useful to produce a better overview of the ontology content, was also created from the regular GO annotation data.

For KEGG pathways annotation, we used Blast2GO to retrieve Enzyme commission (EC) numbers associated with annotated GO terms, and then mapped these ECs to the corresponding KEGG pathways for each FHM protein sequence.

### Gene name annotation

Gene names were assigned based on the top-hit homologous genes from the BLAST search results used for GO annotation. Specifically, for a protein sequence where a top blast hit had an e-value <= 10^−6^ and a Blast2GO annotation score >=55, the gene associated with the protein was assigned the same name, as defined in UniProtKB database (https://www.uniprot.org/uniprot), of the top-hit homologous gene. For a protein sequence with a top BLAST hit e-value <= 10^−6^ but an annotation score <55 or with blast e-value >10^−6^ but <= 10^−5^, the gene was assigned the name of top-hit homologous gene, but “-like” added to the original gene name. If the top hit was already appended with “-like”, it was kept the same, i.e., no “-like -like” gene names were generated; for genes without a blast hit or with hit e-value > 10^−5^, no name was assigned.

### Comparative genomic analysis

Comparative genomic analysis between FHM and zebrafish was primarily conducted using CoGe (https://genomevolution.org/coge/) [81]. The ZF reference genome used for the comparison is the Tuebingen strain (vUCSC, id23058) version, soft-masked by RepeatMasker. The FHM reference genome uploaded to CoGe (ID# id58916) was hard-masked by RepeatMasker. Using repeat-masked version helps to improve and speed up the synteny analysis between the two genomes. The syntenic analyses were performed on CDS and genomic sequences, separately. The default fastest but sensitive aligner LAST (version 744) [82] was selected as the alignment tool for identifying syntenic region. For CDS-based syntenic analysis, the algorithm for merging syntenic blocks and for syntenic depth were both set to be Quota align [83]. For genomic-based syntenic analysis, syntenic path assembly (SPA) [84] analysis was used to create a better visualization of the syntenic map of the two reference genomes. All syntenic plots were generated by SynMap2 [85].

## RESULTS

### De novo assembly of the FHM genome

#### Genome assembly

The whole FHM genome assembling process included eight major steps (Figure 1). In the first step, CANU (version 1.8) was used for base correction, read trimming, and assembly of the initial diploid reference genome from long PacBio reads. Two different sets of long PacBio sequencing data, which include a total 9,361,408 reads with about 75 giga base pairs (Gbps), were generated for the de novo assembly of the FHM genome. As the estimated FHM genome size is 1.1 Gbps [20, 86], the coverage depth of PacBio reads is more than 70X, providing enough depth coverage for the assembly of FHM genome of reasonable quality. The size of the initial draft diploid genome assembly was 1,454,900,621 with a contig N50 (the shortest contig length needed to cover 50% of the genome) of 141,276 bps. In the second step, the diploid draft assembly was polished by Pilon (version 1.23), using 2X100 bps Illumina paired-end reads (SRR1304883 and SRR1301972) and 2-lanes of Illumina 2X250 bps reads generated in-house (see Materials and Methods for more details), to correct base calling errors and fill mini gaps in the draft. Over 99% reads in both Illumina datasets were mapped to the draft assembly, indicating the completeness of the assembly. In the third step, redundant haplotigs, resulting from heterozygous regions of haplotype copies not recognized by the assembler, were purged from the polished draft. This resulted in a haploid assembly of 926,182,690 bps with a contig N50 of 297,600 bps (see Figure 2 and Figure 3). The removal of haplotigs reduced the size of the assembly by almost 520Mbps. In the fourth step, the haplotigs-purged draft was polished using the same sets of llumina reads as used to do the initial polishing. There was little change in the read mapping rate, i.e., remaining over 99%, in polishing run for the assembly before and after purging haplotigs, suggesting only duplicated haplotigs were purged.

**Figure 1:**
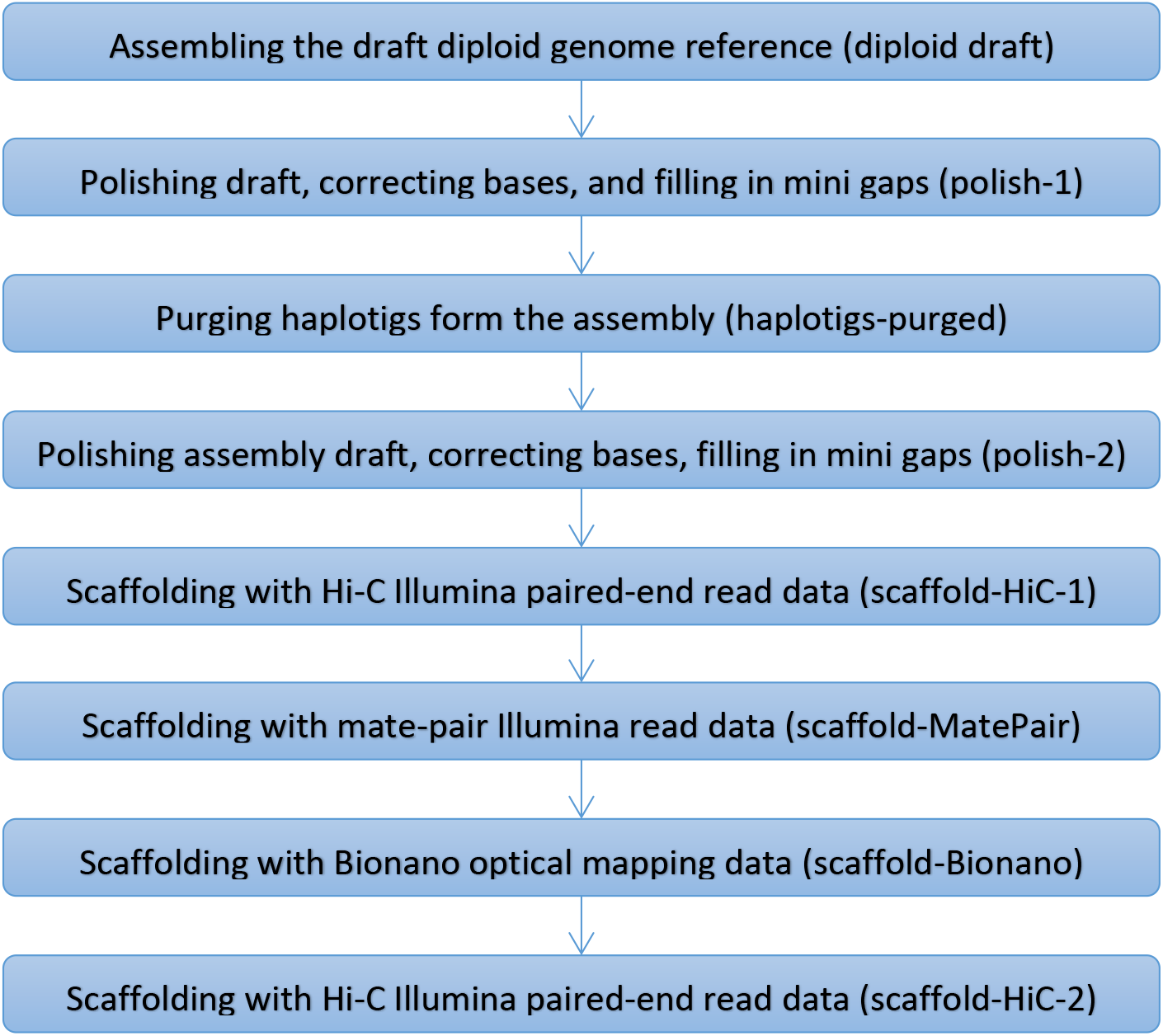
Flowchart for assembling the final FHM2 reference genome. The assembling process includes eight main steps in the order from top to bottom. The assembly steps and order were optimized to achieve the best assembly. For convenience, the brief phrase in parentheses of each step is henceforth used to refer to the corresponding assembly step.

**Figure 2:**
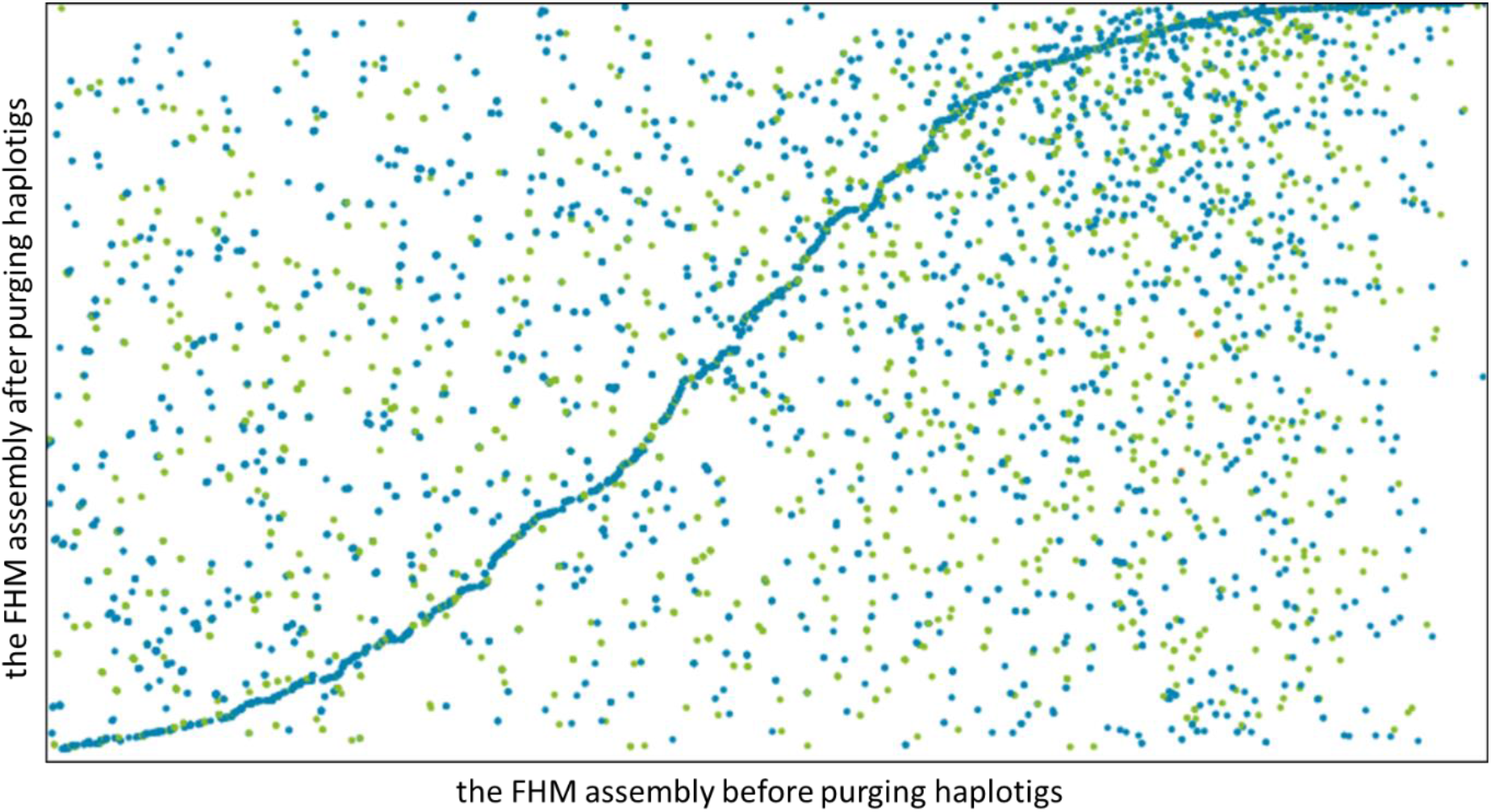
Dotplot of FHM genome assemblies before and after purging haplotigs. The x-axis is the original FHM diploid genome assembly (1.45Gbps) by CANU (version 1.8), and y-axis is the one (0.93Gbs) after purging haplotigs. The removal of haplotigs reduce the size of the FHM genome assembly by almost 520Mbps.

**Figure 3:**
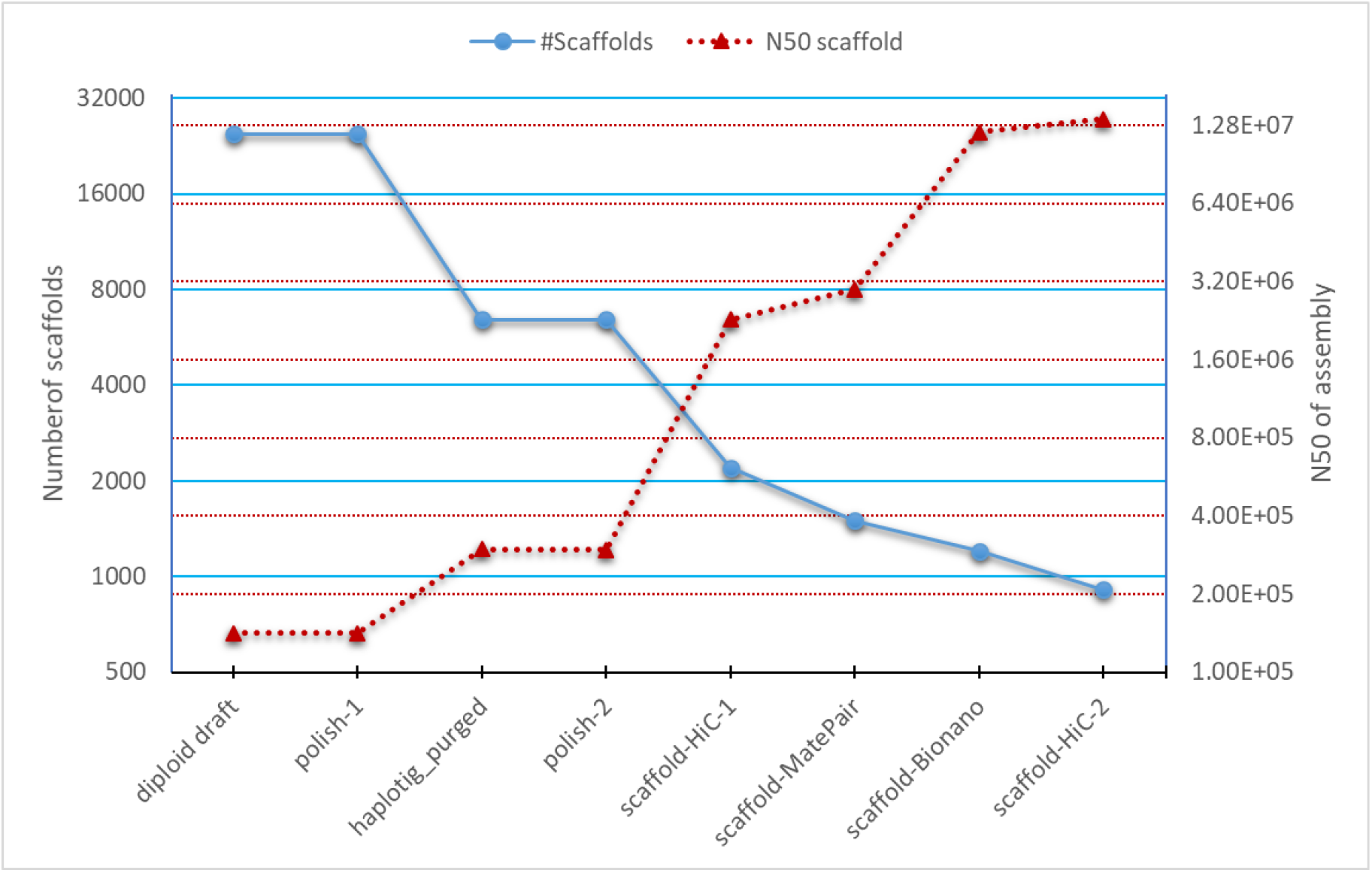
Number of scaffolds and N50 of FHM assemblies at different assembly steps. The x-axis shows different assembly steps from the initial assembling of diploid daft to the final scaffolding with HiC data. The primary y-axis (blue solid line with the round dot marker) is of the number of scaffolds, and the secondary y-axis (red dotted line with triangle marker) is N50. Both Y-axes are in the log2 scale.

To achieve chromosome/sub-chromosome scale assembly, the assembly was then subjected to four rounds of scaffolding (steps 5-8), each increasing scaffold N50 (the shortest scaffold length needed to cover 50% of the genome) of the assembly substantially (Figure 3). The initial scaffolding step with HiC data (14,719X genome overage) drastically increased scaffold N50 of the assembly by eight-fold. Scaffolding with long mate-pair Illumina reads increased N50 only slightly, but it helped to correct some initial scaffolding errors. The incorporation of Bionano data (113X effective coverage) increased scaffold N50 an additional four-fold and helped correct mis-assembled and scaffolded regions (Suppl Figure 2). The final scaffolding with HiC data identified and utilized more long-chromosome interaction sites (Suppl Figure 3), which further improved the assembly to the chromosome/sub-chromosome level with scaffold N50 reaching over 13.5Mbps. After trimming Ns at beginnings and ends of all scaffolds, the scaffold N50 of the final FHM assembly was reduced to 12.0Mpbs. The longest scaffold (over 60Mbps) of the FHM assembly was equivalent in size to the longest chromosome (about 62Mbps) of the closely related zebrafish, suggesting the assembly approaches the chromosome scale.

The final haploid FHM assembly contains 910 scaffolds and 6281 contigs with a total 1,066,412,313 bps in length. Among the 910 scaffolds, 133 (14.6%) of all scaffolds are longer than 1Mbps. The first 15 scaffolds are of chromosome/sub-chromosome size, each >15Mbps, and comprise about 40% the genome sequences (Figure 4, see Suppl Figure 4 for the full distribution). Approximately 90% of the genome is contained on the first 125 scaffolds with greater than 50% on the first 23 scaffolds. The remaining 785 scaffolds accounts for less than 10% genome. For convenience, we henceforth refer our final haploid FHM assembly as FHM2 and the previously published FHM assembly as FHM1.

**Figure 4:**
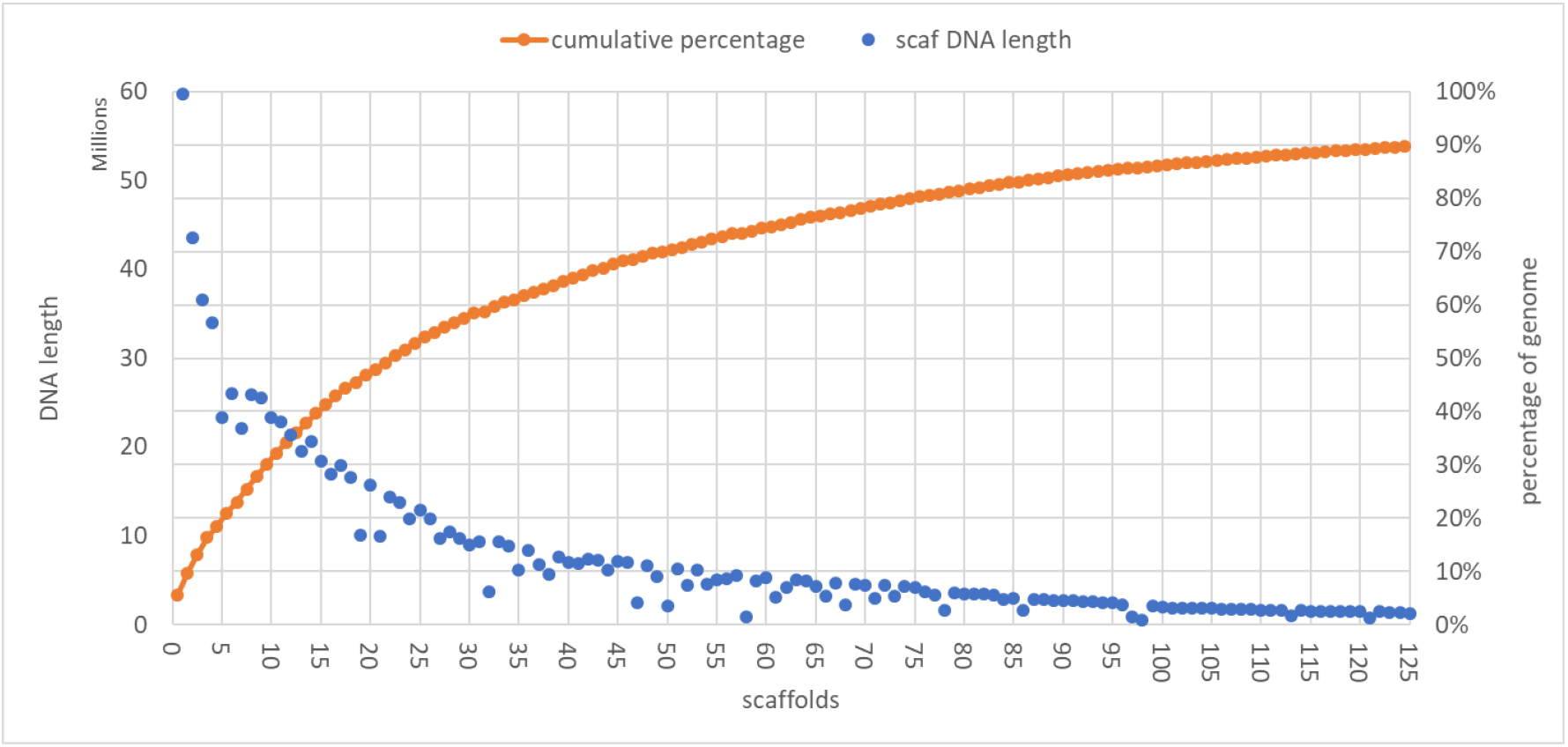
Scaffold length distribution of the FHM genome assembly. The blue dots are lengths of individual scaffolds, and yellow ones are the cumulative percentage of the total genome length. The plot shows only the first 125 scaffolds.

#### Genome assembly assessment

We evaluated the FHM2 assembly in three aspects: 1) contiguity by contig/scaffold N50, the shortest contig/scaffold length in the assembly to cover 50% of the total genome, 2) completeness by BUSCO (Benchmarking Universal Single-Copy Ortholog) score, which checks the presence or absence of highly conserved genes in an assembly, and 3) base accuracy by mapping rates of short Illumina reads to the genome. The contig N50 of the assembly is about 300Kbps and scaffold N50 reaches 12.0Mbps, both of which are much greater than those of FHM1, indicating substantial improvement of FHM2 over FHM1. The BUSCO analysis indicated that the FHM2 assembly is over 95% complete, which approximates the quality of the latest version of the ZF assembly (GRCz11) as shown in Table 1. In addition, the complete and single-copy BUSCOs score of the FHM2 assembly is near 90%, suggesting the high degree of completeness of the FHM2 assembly is coupled with few duplications. The base accuracy of the assembly was assessed indirectly by measuring mapping rate of Illumina genome sequencing reads, assuming a low error rate of bases in assembly lead to a high mapping rate. The overall mapping rates were over 99.4% for both 2X250bps Illumina paired-end datasets that were used to polish the assembly, indicating a low base error rate of the assembly.

**Table 1:** Comparison of three genome assemblies. FHM1 is the first FHM genome assembly currently publicly available. ZF assembly (GRCz11 primary assembly) is the latest version available at the NCBI and the Ensembl.

| Assembly statistics | FHM1<br>(GCA_000700825) | FHM2<br>(WIOS000000000) | ZF<br>(GRCz11) |
| --- | --- | --- | --- |
| Number of scaffolds | 73,057 | 910 | 993 |
| N50 contig | 7,513 | 300,151 | 1,428,257 |
| N50 scaffold | 60,380 | 11,952,773 | 54,304,671 |
| Complete BUSCOs | 3,506 (76.5%) | 4,357 (95.1%) | 4,384 (95.7%) |
| Complete and single-copy BUSCOs | 3,324 (72.5%) | 4,115 (89.8%) | 4,215 (92.0%) |
| Complete and duplicated BUSCOs | 182 (4.0%) | 242 (5.3%) | 169 (3.7%) |
| Fragmented BUSCOs | 507 (11.1%) | 73 (1.6%) | 66 (1.4%) |
| Missing BUSCOs | 571 (12.4%) | 153 (3.3%) | 134 (2.9%) |
| Total contig size | 811,183,656 | 925,375,343 | 1,368,782,359 |
| Total scaffold size | 1,219,326,373 | 1,066,412,313 | 1,373,471,384 |

### Repeat annotation

We used RepeatModeler (version 2.0.1) and RepeatMasker (version 4.1.0) to annotate repetitive regions of the FHM2 genome assembly (see Methods for the details). Overall, we found that more than 461Mbp genome sequences, or 43.27% of the FHM2 genome assembly, were repetitive regions (Table 2). Among all repeat elements, DNA transposons were the most abundant, accounting for 21.47% of the whole genome. In contrast, only 9.09% of the genome were of retroelements, which were dominated by long terminal repeats (LTR) that represented about 6.62% of the genome.

**Table 2:** Repeats analysis summary report of the FHM2 genome assembly

| Type | sub-group | # elements* | length (bp) | % genome |
| --- | --- | --- | --- | --- |
| **Retroelements** |  | 249678 | 96792759 | 9.08% |
| **SINEs** |  | 22354 | 2714953 | 0.25% |
|  | Penelope | 1207 | 265815 | 0.02% |
| **LINEs** |  | 78893 | 23445138 | 2.20% |
|  | L2/CR1/Rex | 59637 | 15291306 | 1.43% |
|  | R1/LOA/Jockey | 2299 | 857151 | 0.08% |
|  | R2/R4/NeSL | 2263 | 1224253 | 0.11% |
|  | RTE/Bov-B | 4905 | 1780570 | 0.17% |
|  | L1/CIN4 | 2184 | 1032311 | 0.10% |
| **LTR elements** |  | 148431 | 70632668 | 6.62% |
|  | BEL/Pao | 9750 | 6260165 | 0.59% |
|  | Ty1/Copia | 406 | 234805 | 0.02% |
|  | Gypsy/DIRS1 | 89802 | 47549943 | 4.46% |
|  | Retroviral | 17549 | 7893150 | 0.74% |
| **DNA transposons** |  | 1138712 | 228911213 | 21.47% |
| **hobo-Activator** |  | 256864 | 54737880 | 5.13% |
| **Tc1-IS630-Pogo** |  | 71534 | 14918781 | 1.40% |
| **PiggyBac** |  | 19182 | 5854328 | 0.55% |
| **Tourist/Harbinger** |  | 89007 | 19615220 | 1.84% |
| **Other (Mirage,P-element, Transib)** |  | 8307 | 2261427 | 0.21% |
| **Rolling-circles** |  | 43452 | 22236361 | 2.09% |
| **Unclassified** |  | 583472 | 89130569 | 8.36% |
| **Total interspersed repeats** |  |  | 414834541 | 38.90% |
| **Small RNA** |  | 3015 | 513354 | 0.05% |
| **Satellites** |  | 30422 | 5491099 | 0.51% |
| **Simple repeats** |  | 354572 | 16612360 | 1.56% |
| **Low complexity** |  | 35283 | 1920018 | 0.18% |

\*most repeats fragmented by insertions or deletions have been counted as one element

*We compared the repeat distribution of the FHM2 assembly to those of repeats statistics [87] reported for four other teleost genomes, zebrafish, medaka (*Oryzias latipes*), stickleback (*Gasterosteus aculeatus*), and pufferfish (*Tetraodon nigroviridis*). The five teleost species differ substantially in terms of the distribution of repeat elements (Suppl Table 4*). The zebrafish genome has the highest proportion (56.49%) of total interspersed repeats, while the FHM genome is the second highest (38.90%). In contrast, the AT-rich FHM genome contains the highest proportion of simple repeats and low complexity regions among all five species.

### Gene annotation

#### Gene prediction

The ab initio gene prediction by AUGUSTUS (Version 3.3.3), conducted without using any RNA-seq or EST data as evidence, identified a total of 251,638 coding exons and 26,985 genes in the FHM2 assembly. The BUSCO analysis of the resulting proteins showed 95.6% BUSCO genes were present, but 10.7% were fragmented, indicating that the predicted genes for the FHM2 were relatively complete. In order to improve gene prediction further, a pipeline consisting of the popular gene prediction tools Maker and EVidenceModeler (EVM) was developed for predicting gene models with RNA-seq data (Suppl Figure 5).

Maker produced 30,909 gene models with 47,716 transcripts/proteins, and the BUSCO analysis of its exemplar protein set, which contains only the longest protein from each gene, showed the complete BUSCO score at 84.8% with 5.8% duplication. RNA-seq read mapping analysis reported the overall mapping rate to Maker’s exemplar transcript set (transcripts corresponding to the longest proteins of each gene) was 77.9% and unique mapping rate was 71.7%. Performed better than Maker, EVM predicted a total of 37,190 gene models, of which the exemplar transcripts had a complete BUSCO score of 94.1% with 6.7% duplication. The total RNA-seq mapping rate to EVM’s exemplar transcripts was 77.9% with the unique mapping rate at 72.9%. To refine EVM gene models with UTR information, the 37,378 EVM gene models were run back through Maker. 36,881 gene models were retained, including those EVM models that have no overlapping coding sequences (CDS) with the Maker 30,909 gene models. The new set of 36,881 genes had a complete BUSCO score of 94.2% with 6.6% duplication, and the overall RNA-seq read mapping rate of 85.3% with 78.5% unique mapping rate, which were better than either of the original EVM or Maker sets.

We applied a filter, based on mapping evidence and homology to reference proteins (see gene models filtering section in Methods) to the 36,881 gene models to remove potential pseudo genes or non-functional paralogs. After filtering, the number of remaining gene models was 26,150 with 47,578 transcripts/proteins. Little change in either complete BUSCO score or overall read mapping rate was observed. The filtered gene models were then manually curated to generate the final set of gene models, transcripts, and protein sequences. In our manual curation, if an original EVM protein sequence’s best BLAST [88] hit to a reference protein was better than that of the corresponding protein from the final Maker run, the original EVM protein was returned to the final models, replacing the protein that the final Maker run had modified. 303 of the total 47,578 protein sequences were changed back to their original EVM sequences based on this criterion. Most protein changes were due to different translation of existing transcripts. Only six transcripts were changed in the manual curation. One to six bases were either added or removed from five of those six transcripts while one exon of 206 bases was added to the last one. The final curated gene set has a complete BUSCO score of 94.3%, slightly improved compared to pre-curation (Suppl Figure 1), while maintaining the read mapping rate at 84.9% (Suppl Table 5).

#### Benchmark comparison with zebrafish

In order to determine the quality of the FHM gene models, a benchmark analyses were carried out in comparison with the zebrafish, which is closely related to the FHM and has the best characterized gene models in fish species. For this comparison, the full set of protein or transcript sequences rather than the exemplar set were used for both BUSCO and read mapping rate analysis.

The results show comparable overall BUSCO scores between FHM and ZF (Table 3). The complete BUSCO score for FHM proteins is 94.3% and for FHM transcripts is 94.7%, which are only slightly lower the corresponding score for ZF. However, the complete and duplicated BUSCO scores for FHM are noticeably lower (8.2% and 6.1% lower for proteins and transcripts, respectively) than those for ZF, suggesting our FHM gene models potentially have a lower duplication level.

**Table 3:**
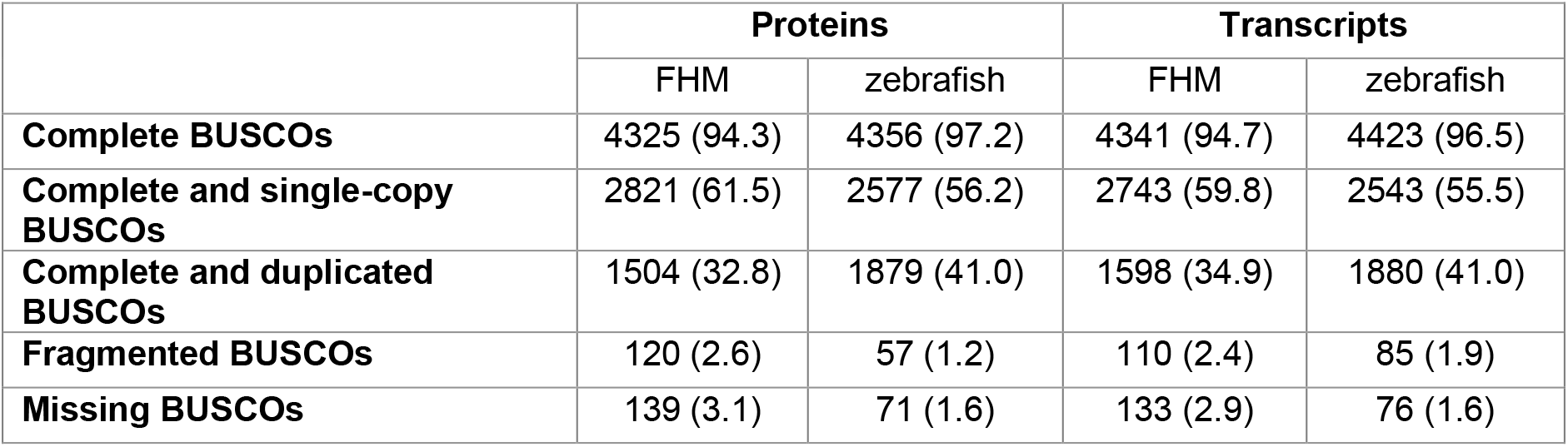
Comparison of protein and transcript BUSCO scores between FHM and ZF. The numbers in parentheses are the corresponding percentages of the BUSCOs.

|  | Proteins |  | Transcripts |  |
| --- | --- | --- | --- | --- |
|  | FHM | zebrafish | FHM | zebrafish |
| **Complete BUSCOs** | 4325 (94.3) | 4356 (97.2) | 4341 (94.7) | 4423 (96.5) |
| **Complete and single-copy BUSCOs** | 2821 (61.5) | 2577 (56.2) | 2743 (59.8) | 2543 (55.5) |
| **Complete and duplicated BUSCOs** | 1504 (32.8) | 1879 (41.0) | 1598 (34.9) | 1880 (41.0) |
| **Fragmented BUSCOs** | 120 (2.6) | 57 (1.2) | 110 (2.4) | 85 (1.9) |
| **Missing BUSCOs** | 139 (3.1) | 71 (1.6) | 133 (2.9) | 76 (1.6) |

The read mapping analysis shows similar overall mapping rates of 50bp RNA-seq reads (Table 4). The transcriptome mapping rates for FHM are all slightly better than those for ZF. In particular, the unique transcriptome read mapping rates for FHM are 4-5% higher than the corresponding ones for ZF. Whole genome mapping rates were consistent, though slightly less pronounced, i.e., about ~3% difference, with transcriptome read mapping results. Together, BUSCO and mapping rate analysis indicates that the FHM2 genome assembly and gene prediction models are of at least similar quality to ZF.

**Table 4:** Comparison of RNA-seq read mapping rates between FHM and zebrafish. To make comparison fair, zebrafish primary assembly (GRCz11 primary assembly) was used and only protein transcript sequences from the primary reference were used. Reported values in each cell are total mapping rate, unique mapping rate, multimapping rate, respectively

|  | WHOLE GENOME |  | TRANSCRIPTOME |  |
| --- | --- | --- | --- | --- |
|  | FHM | Zebrafish | FHM | Zebrafish |
| **BOTH READS** | 90% (80%, 10%) | 87% (79%, 8%) | 74% (45%, 29%) | 69% (39%, 30%) |
| **1<sup>ST</sup> READ** | 94% (81%, 13%) | 91% (80%, 11%) | 81% (45%, 36%) | 75% (40%, 35%) |
| **2<sup>ND</sup> READ** | 92% (80%, 12%) | 90% (79%, 11%) | 78% (45%, 33%) | 75% (40%, 35%) |

The number of predicted protein coding genes and transcripts in the FHM2 genome is very close to that in the ZF genome, which has 25,638 protein genes and 47,834 corresponding transcripts (Table 5). The two species are also similar in the number of transcript isoforms per gene (1.87 for ZF and 1.82 for FHM). There are, however, some noticeable differences. First, the proportion of transcripts without 5’ or 3’ UTR in FHM2 is over 18%, which is far greater than about 5% in ZF. This large difference could reflect incomplete UTR prediction in FHM rather than, the actual biological differences between the two species. Second, the median protein coding gene length in FHM2 is 10,177bps, much shorter than 13,569bps in ZF. The gene length difference is likely to due to more compact gene structure in the FHM2 genome, which is further supported by the observation that the median intron length in FHM2 (469bps) is also much shorter than in ZF (984bps).

**Table 5:**
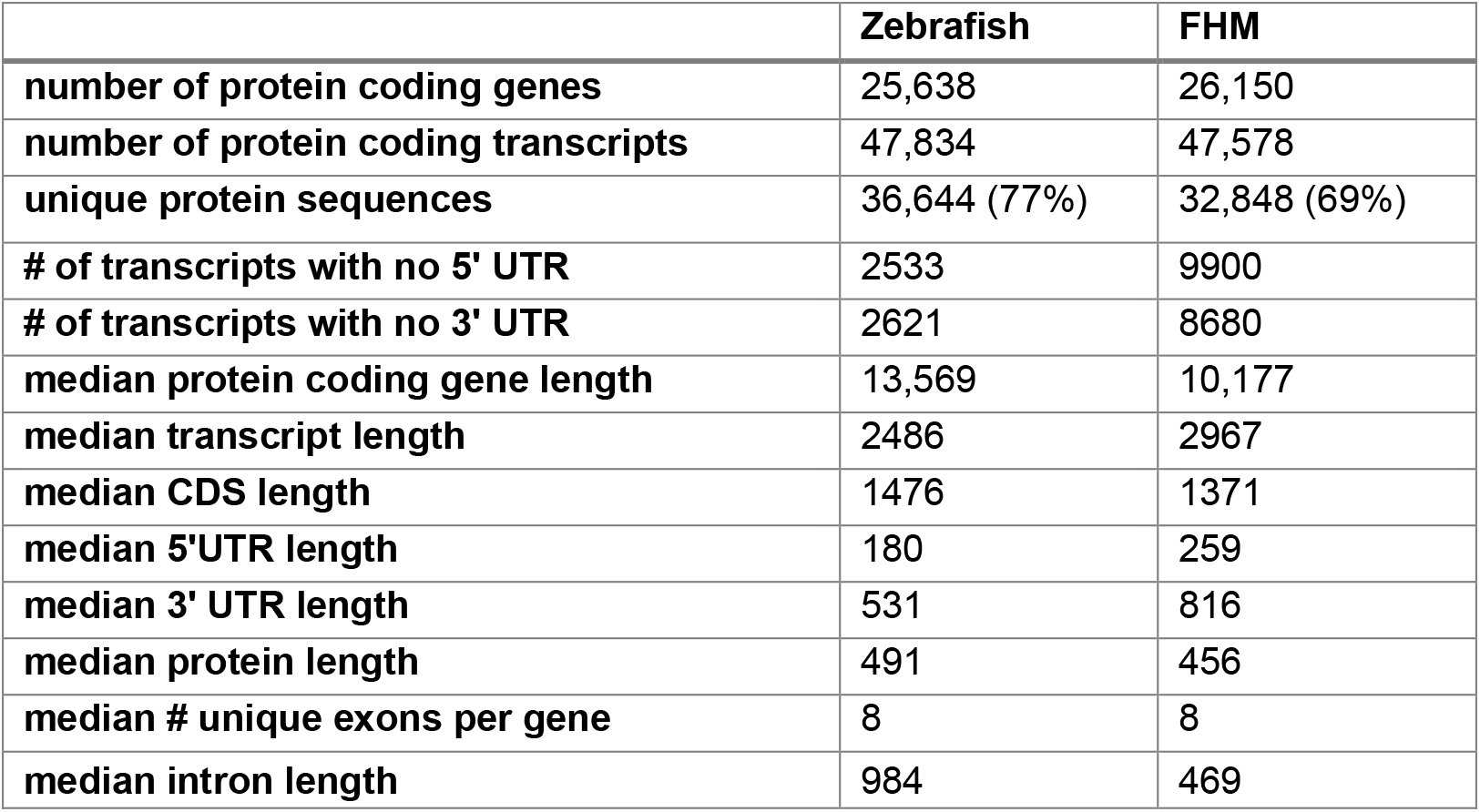
Comparison of FHM model transcripts/proteins to transcripts/proteins extracted from primary assembly of Danio rerio (GCF_000002035.6_GRCz11)

|  | **Zebrafish** | **FHM** |
| --- | --- | --- |
| **number of protein coding genes** | 25,638 | 26,150 |
| **number of protein coding transcripts** | 47,834 | 47,578 |
| **unique protein sequences** | 36,644 (77%) | 32,848 (69%) |
| **# of transcripts with no 5' UTR** | 2533 | 9900 |
| **# of transcripts with no 3' UTR** | 2621 | 8680 |
| **median protein coding gene length** | 13,569 | 10,177 |
| **median transcript length** | 2486 | 2967 |
| **median CDS length** | 1476 | 1371 |
| **median 5'UTR length** | 180 | 259 |
| **median 3' UTR length** | 531 | 816 |
| **median protein length** | 491 | 456 |
| **median # unique exons per gene** | 8 | 8 |
| **median intron length** | 984 | 469 |

#### Gene name and functional annotation

Gene names and functions of predicted genes were annotated through BLAST-searching homologs in reference species and through InterProScan-searching functional domain signatures against the InterPro database. In the BLAST search against the 27 species in the NCBI landmark database, zebrafish (*Danio rerio*) was the species returning the highest number of top BLAST hits (Figure 5); 93% of predicted FHM2 transcripts had a top-matched homologous gene in the ZF genome, confirming the species high degree of evolutionary relatedness. InterProScan signature search also successfully found matched functional domain signatures in the InterPro database for 96% of all predicted FHM2 transcripts. Overall, 22,743 (87%) among the all 26,150 predicted FHM protein coding genes were assigned a gene name, and 42,940 (90%) transcripts representing 22,469 genes (86%) were annotated with at least one GO term. Two versions of GO annotation were provided: the regular full GO version including all GO terms and the GO slim version with a reduced number of GO terms. The full GO version provides details about the function of a gene while the GO slim version is better for the overview functional enrichment analysis. The full GO version has 873,774 GO terms while the GO slim version has only 498,616 terms. Figure 6 shows the comparison of number of GO terms in three GO categories: biological process (BP), cellular component (CC), and molecular function (MF). In the GO slim version, the largest number of GO terms assigned to a transcript is 50. *Figure 7* shows the detailed distribution of GO terms in the GO slim version among all predicted FHM transcripts.

**Figure 5:**
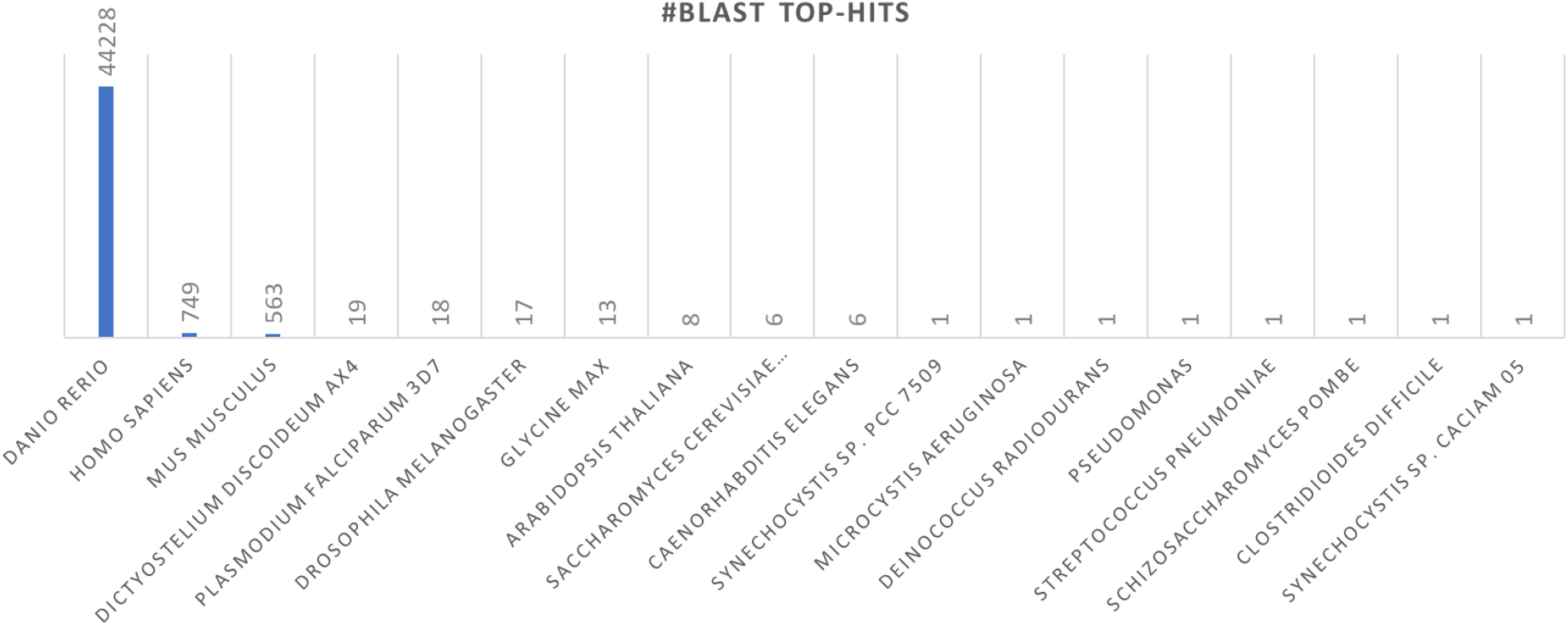
Distribution of 27 landmark species with respect to the number of top-blast hits in homolog search.

**Figure 6:**
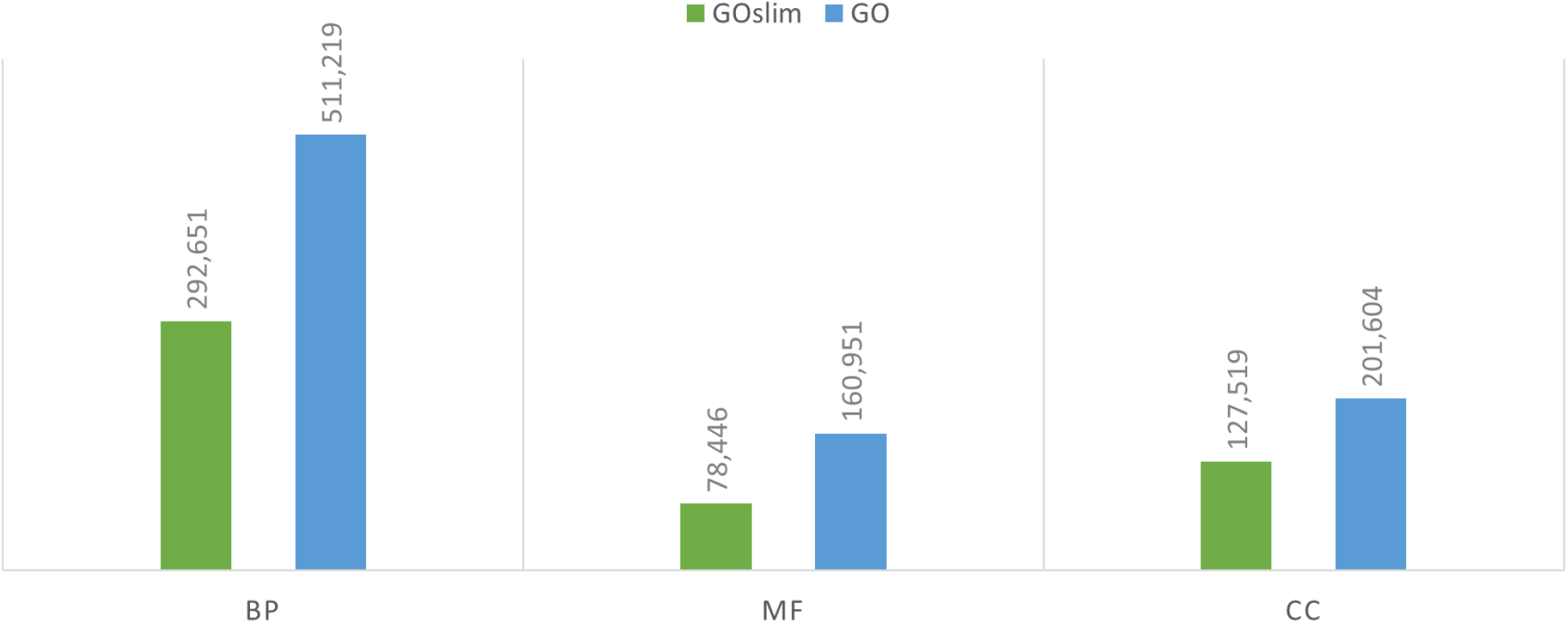
Number of annotated GO terms in BP (Biological Process), MF (Molecular Function) and CC (Cellular Compartment) categories.

**Figure 7:**
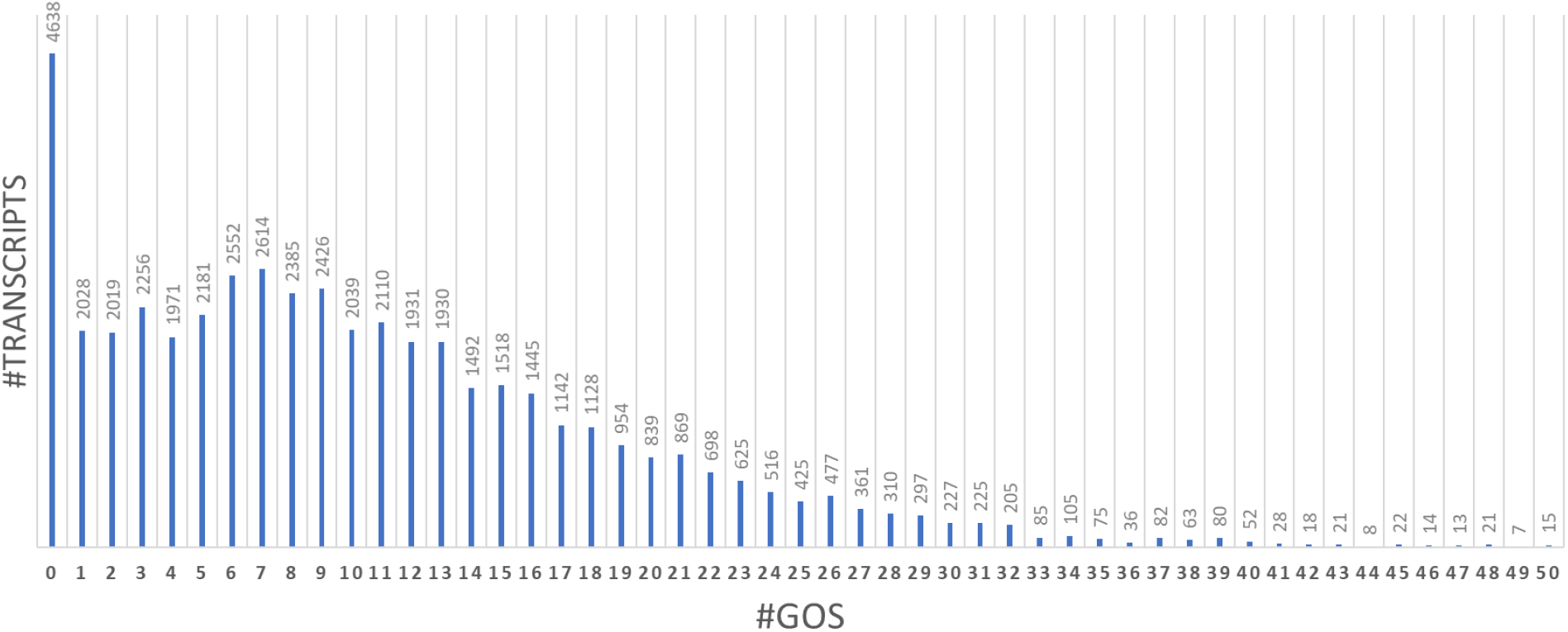
GO term distribution among all predicted transcripts in the FHM2 genome.

In addition, our KEGG pathway annotation identified 148 KEGG metabolic pathways, in which 8,166 transcripts were mapped to at least one KEGG pathway. Furthermore, our annotation found 50,800 enzyme-transcript mapping pairs including 1,401 unique enzyme commission (EC) numbers associated with 12,176 different transcripts.

### Prediction of short non-coding RNAs

#### tRNA prediction

A total of 2670 tRNAs, as a “high-confidence” set, were predicted in the FHM genome by tRNAscan-SE (v. 2.0.5)[74] using the EukHighConfidenceFilter (Table 6, Suppl Table 6). The high confidence set was generated from the initial prediction set with 3776 tRNAs after filtering out pseudogenes and potentially false hits. The high confidence set of tRNAs are most likely to be functional in the translation process [74]. The detailed list of FHM tRNAs by anticodon counts is given in Suppl Table 7, which can be used as a reference for codon usage bias analysis.

**Table 6:** Numbers of different small RNAs predicted in FHM in comparison to reported values for zebrafish

| **Type** | **FHM** | **Danio rerio** | **Source** |
| --- | --- | --- | --- |
| tRNA | 2670 | 8676 | gtRNAdb |
| miRNA | 620 | 387/374 | rfam.xfam.org/mirbase.org |
| piRNA | 392,936 | 1,330,692 | regulatoryrna.org |
| SSU_rRNA_eukarya | 47 | 36 | rfam.xfam.org |
| LSU_rRNA_eukarya | 16 | 52 | rfam.xfam.org |
| snoRNA | 201 | 283 | rfam.xfam.org |
| scaRNA | 13 | 11 | rfam.xfam.org |

#### miRNA prediction

A total of 620 miRNAs were predicted in the FHM genome by miRDeep* using a stringent cutoff of 100. This cutoff was chosen to minimize false positive hits while retaining most predicted mRNAs with exact matched ones in the zebrafish. BLAST search against the total miRNA database at miRbase.org found at least one significant match for 618 of the 620 predicted miRNAs, suggesting most predicted miRNAs are likely to be functional. Compared with the other species, FHM has more miRNAs than most fish species, e.g. zebrafish with only 374 miRNAs, except *Oreochromis niloticus*, however, the mammal species tend to have a far greater number of miRNAs than FHM (Suppl Table 8). The abundance of miRNAs in the FHM genome suggest that post-transcriptional regulation including RNA silencing may play an important role in gene expression regulation in the FHM.

#### piRNA prediction

Piwi interacting RNAs (piRNAs) were also identified in the FHM genome. piRNAs are small non-coding RNAs of length about 24–32 nucleotides. piRNAs typically play regulatory roles by binding to members of the PIWI proteins, and they can also regulate signaling pathways at the transcriptional or post-transcriptional level. In addition, piRNAs and PIWI proteins may be used as biomarkers for a various of cancers as they also often abnormally expressed in cancer tissues. Prediction of piRNAs were accomplished using both Piano [76] and piRNN tools [77]. In the FHM genome, 87,498 transposon-interacting piRNAs were predicted by Piano (see Suppl Figure 11 for length distribution), and a total of 392,936 all piRNAs of length between 25-40 bps were predicted by piRNN (Suppl Figure 12). Based on the prediction, FHM has far fewer piRNAs than ZF, which has about 1.33 million piRNAs in piRBase (http://www.regulatoryrna.org). The abundance level of piRNAs in FHM is nevertheless similar to that (> 300,000 piRNAs) of many other species reported in the study by Ozata et al [89].

#### Other small RNAs prediction

The Infernal/Rfam (IR) programs were used for prediction of other non-coding RNA genes, including rRNA, snRNA, snoRNA, small Cajal body-specific RNA (SCARNA), RNase_MRP), nuclear RNAse P (RNaseP_nuc), and signal recognition particle RNA (Metazo_SRP)(Table 6 and *Suppl Table 9*).

### Data visualization with the UCSC genome browser

We took the advantage of the powerful data visualization functions of the UCSC genome browser for visualization of the FHM2 reference genome and annotation. We built a fully functional FHM assembly hub for the UCSC genome browser (Figure 8). Users can use the assembly hub to visually explore the FHM2 genome reference and annotation with their own experimental data by adding customized browser tracks of data. Such data visualization with the UCSC browser is a critical and popular approach for manually validating analysis results of large sequencing data and/or generating new hypotheses. By using our UCSC assembly hub, the UCSC genome browser can generate a detailed view of gene annotation including both ab initio gene prediction track and the final gene annotation track, searchable by gene symbol or gene accession number. In addition, it has the repeat annotation track, allowing to explore repetitive regions. Our UCSC assembly hub data hosted at CyVerse Discovery Environment (https://de.cyverse.org/de/) are available at (https://data.cyverse.org/dav-anon/iplant/home/myepa/FHM/FHM2assembly_hub.txt).

**Figure 8:**
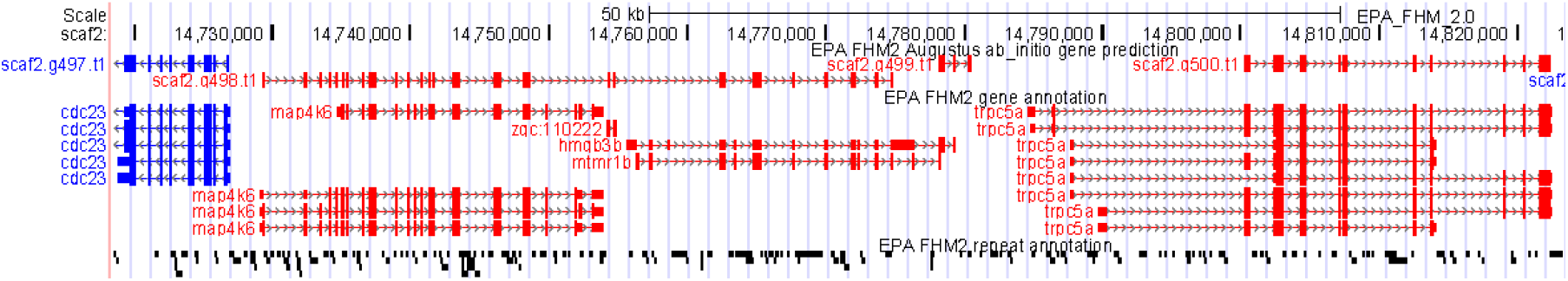
Visualization of FHM2 genome annotation on UCSC genome browser. Tracks are as follows from top to bottom: ab initio gene prediction track; final gene annotation track; the repeat annotation tract shown in the dense mode. Both gene annotation tracks are displayed gene model in the pack mode; genes in red are in the plus strand while those in blue are minus strand of DNA.

Data visualizations with the FHM2 reference genome is also available on CoGe with the EPIC-CoGe browser (https://genomevolution.org/coge/GenomeView.pl) [90]. The EPIC-CoGe brower allows users to visually explore GC content of the reference genome and generate a bird’s-eye view of gene annotation as shown in Figure 9.

**Figure 9:**
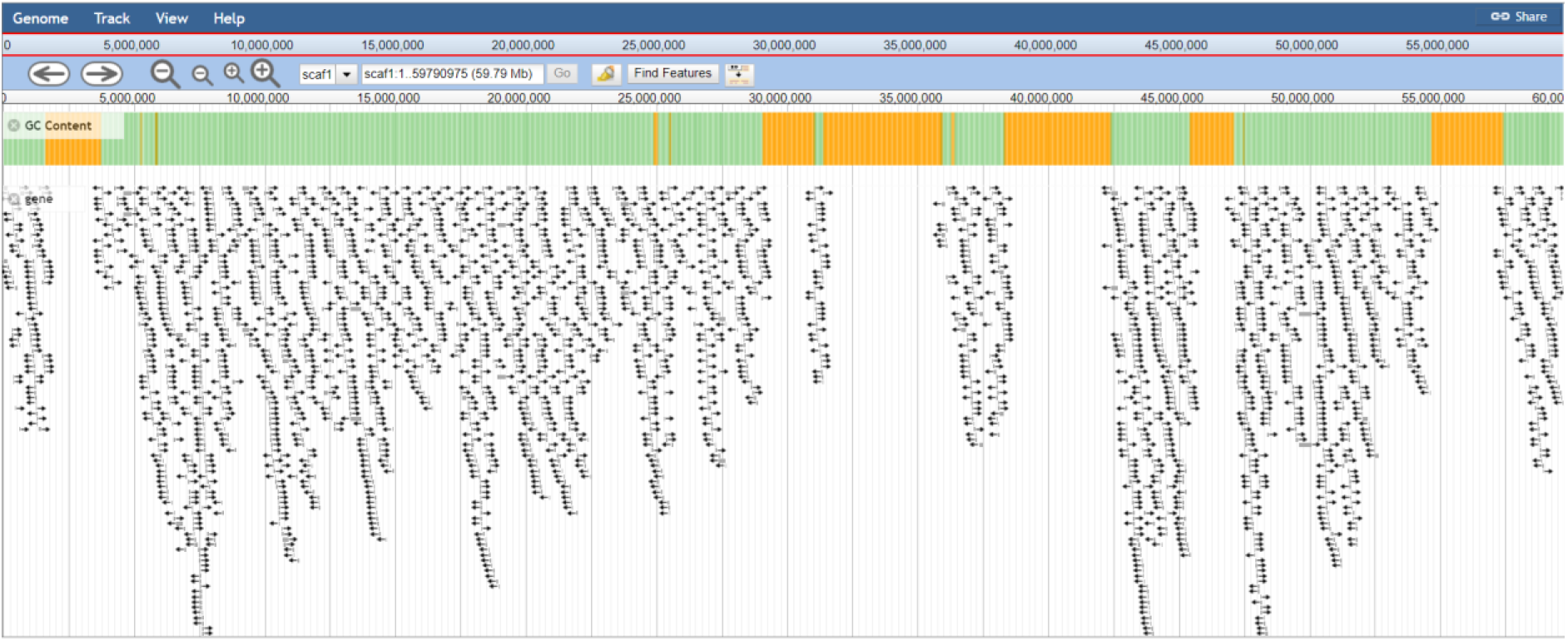
EPIC-CoGe genome browser’s bird’s-eye view of the first scaffold of the FHM2 genome assembly. The top dense color bar indicates GC content, and yellow regions with 0%GC are repeat-masked or gapped region. The arrowed lines are of annotated genes with arrow pointing the direction of genes.

### Comparison with the Zebrafish genome

#### Synteny analysis

The comparison analysis of the ZF and FHM genome reveals that they share extensive syntenic regions. In some cases, syntenic regions are almost of the same length of an entire scaffold or chromosome as shown by syntenic dot-plots (Figure 10 and Suppl Figure 6). The CDS-based syntenic analysis shows there are 117,846 high-scoring segment pairs (HSPs) from the LAST [82] alignment after filtering out tandem duplicates. The total length (74,989,746) of HSPs accounts for over 7% or 8% if excluding scaffolding gaps) of the FHM2 reference genome. The average length of HSPs is 636 bps with the mean percent identity at 71% the maximum at 31,479bp (Suppl Figure 10). Such large syntenic regions shared between the two species indicates the two species are closely related evolutionarily. Upon closer inspection of these syntenic regions using the GEvo tool, however, we found that genomic regions containing syntenic blocks are substantially different between two species. Particularly, genomic regions in the FHM genome are often much shorter than the corresponding regions in the ZF genome. In most syntenic genomic regions that we examined with the GEvo tool, the length of an FHM region is only about one half of the corresponding ZF region. For example, the plot of syntenic blocks (Figure 11) shows that the genomic region surrounding the transcript FMt022379 in FHM is only about 90kbps in length while the corresponding region surrounding the transcript ENDART00000132491 (gene ENSDARG00000025554) in ZF is 200kbps, more than twice as long (see Suppl Figure 8 and Suppl Figure 9 for additional examples). Such difference in length can be also seen in the syntenic dot-plot shown in Figure 10, where lengths of FHM syntenic regions represented by the y-axis are much shorter than those of the corresponding ZF represented by the x-axis. The results suggest that FHM and ZF have diverged substantially from their last common ancestor, and ZF may have gained much more intergenic or intron regions than FHM, which has either gained little or lost such non-coding regions in the evolutionary process from their ancestor. This also indicates that FHM has a much more condensed genome than ZF as the two species have a similar number of genes.

**Figure 10:**
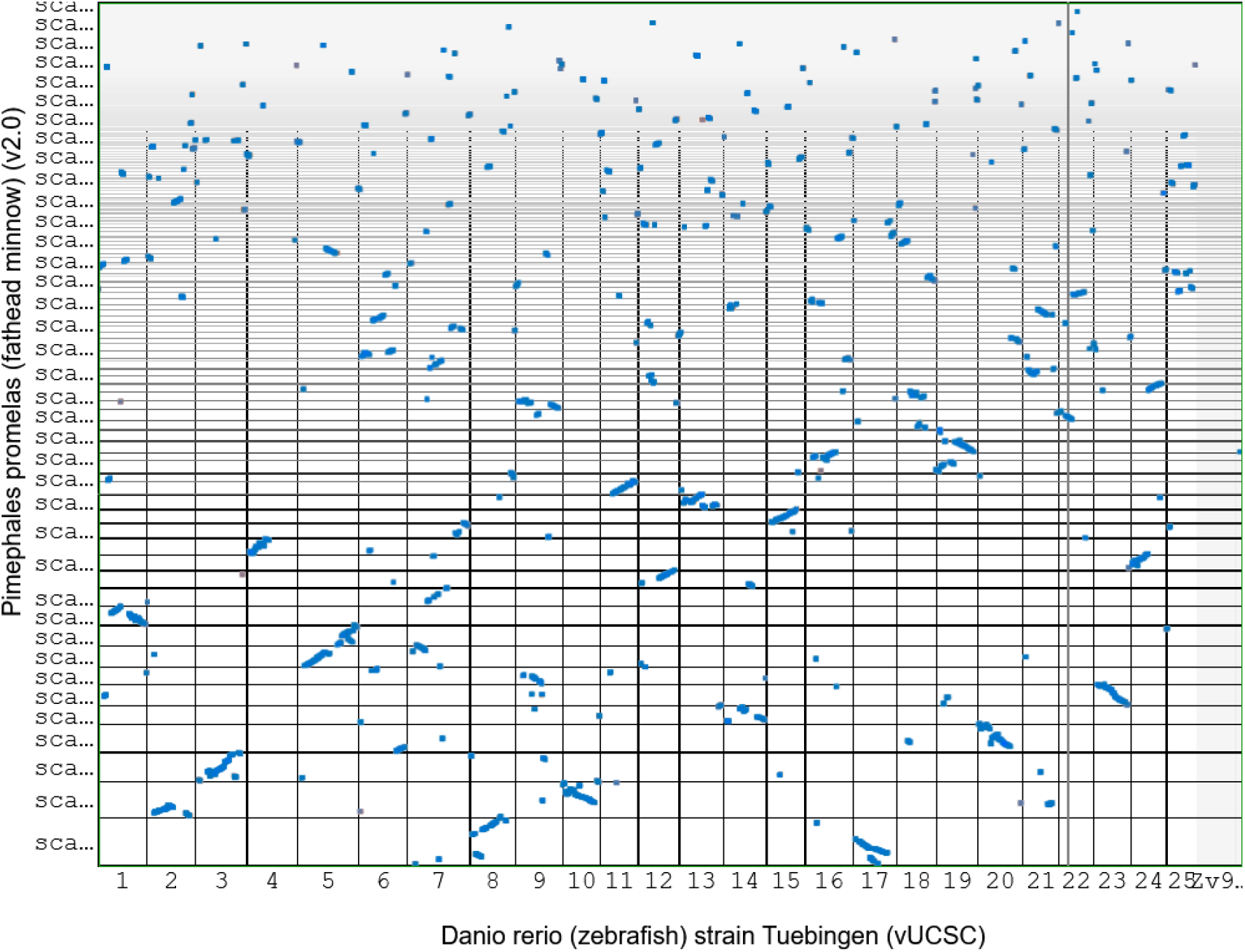
Syntenic dot-plot of CDS regions of FHM and Zebrafish. x-axis is zebrafish and y-axis is FHM. The plot was generated by SynMap2.

**Figure 11:**
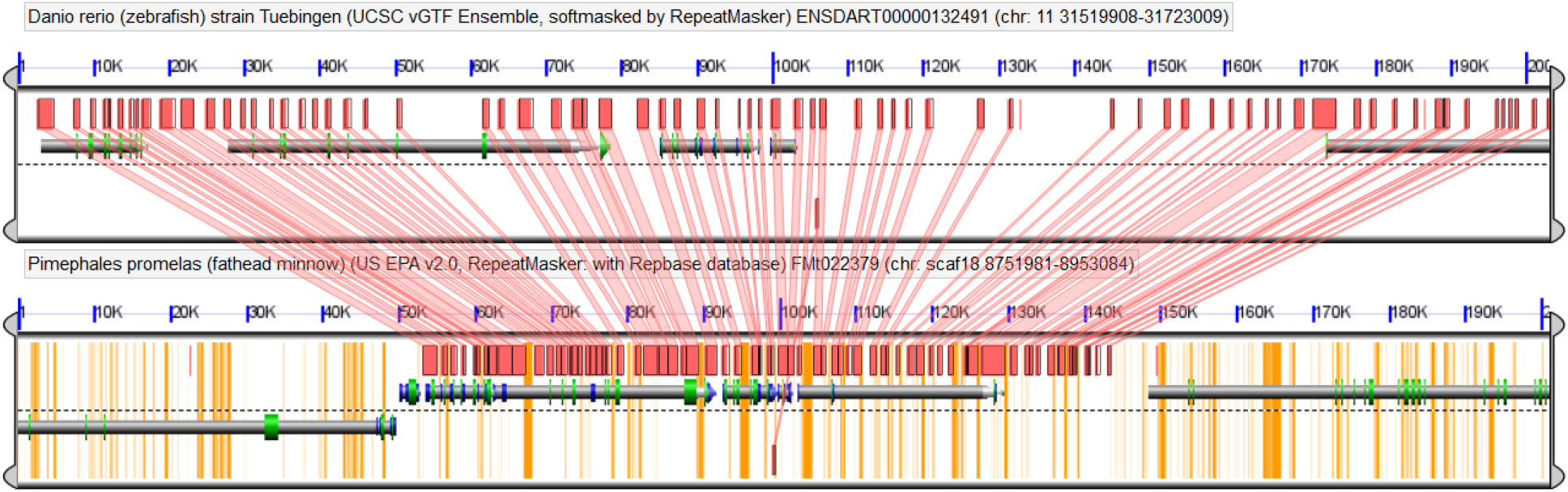
GEvo syntenic map of the genomic region surrounding the FMt022379 in FHM2 and the ENDART00000132491 in zebrafish. The genomic region containing the syntenic region in the ZF is approximately 200kbps, more than twice the length of the corresponding region (about 90kbps) in the FHM2 genome. In this syntenic region, the FHM genome has much shorter introns or intergenic regions. The yellow regions in the FHM genome are repeat-masked or gap regions.

The synteny analysis also revealed the mutation rates of syntenic regions between the two species with the synonymous substitution rate (Ks) being 0.62 and the nonsynonymous substitution rate (Ka) being 0.10 (see Suppl Figure 7). The Ka/Ks ratio (0.16) between FHM and ZF is far less than 1, indicating strong purifying/negative selection pressure in their evolutionary process.

#### Conservation of estrogen response and exposure biomarker genes

FHM and ZF are among the most widely used model fish species in environmental toxicogenomics. In particular, toxicogenomic approaches have been applied to the endocrine-disrupting chemicals (EDCs), e.g.,17α-ethynylestradiol (EE2) to better characterize the cellular responses underlying their adverse effects on reproduction and development [1]. Many studies have measured the differential expression of known estrogen responsive genes across species as a means of determining estrogenic exposure, however, these studies have largely been done without consideration of the underlying genomic structure. For this reason, we selected two well-studied estrogen-responsive genes, estrogen receptor 1 gene (ESR1), a critical gene for regulating estrogen response and the vitellogenin gene (VTG), an egg yolk precursor and a biomarker indicating exposure to estrogen-like chemicals in male fish, to determine the degree of conservation between the two species. Both FHM and zebrafish genomes contain a single copy of the ESR1 gene, however, eight transcript isoforms were identified in the FHM compared to four in ZF. The syntenic alignment of the ESR1 genes shows they are highly conserved in exon order and all CDS or CDS flanking region sequences, but differ markedly in their intron regions with large deletions and insertions (Figure 12). In contrast to ESR1, seven copies of the VTG gene are found in both species. Both their sequence and genomic arrangement appears to be highly conserved between the two species. In both species, six of the seven VTG genes are tandemly arrayed within a tight region of a single chromosome or scaffold (chr22 in ZF, scaf140 in FHM)(Figure 13). The high copy number is likely the result of gene duplication events in the evolutionary process. Another VTG gene, VTG3, is in a different chromosome, chr11 and scaf62 in ZF and FHM, respectively. Sequences of the ortholog VTG genes are highly conserved in all exons. Conservation of intron sequences was also observed except for the largest intron in the ZF, which is greatly reduced in the FHM (Figure 14).

**Figure 12:**
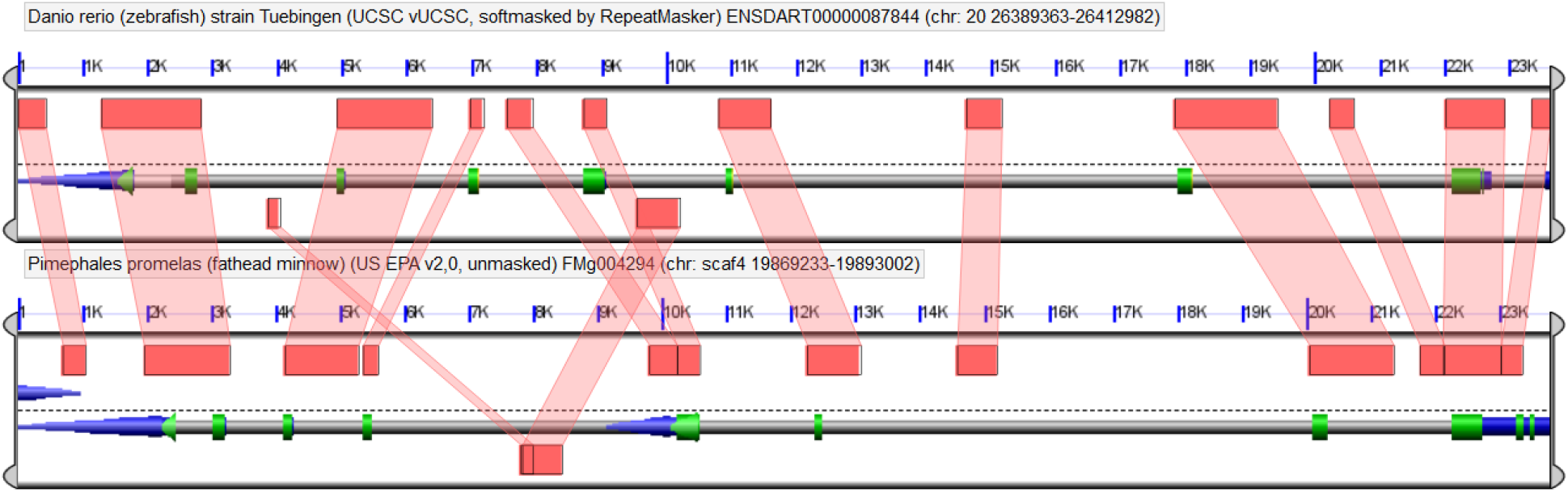
Syntenic alignment map of the ESR1 genes between zebrafish and FHM2

**Figure 13:**
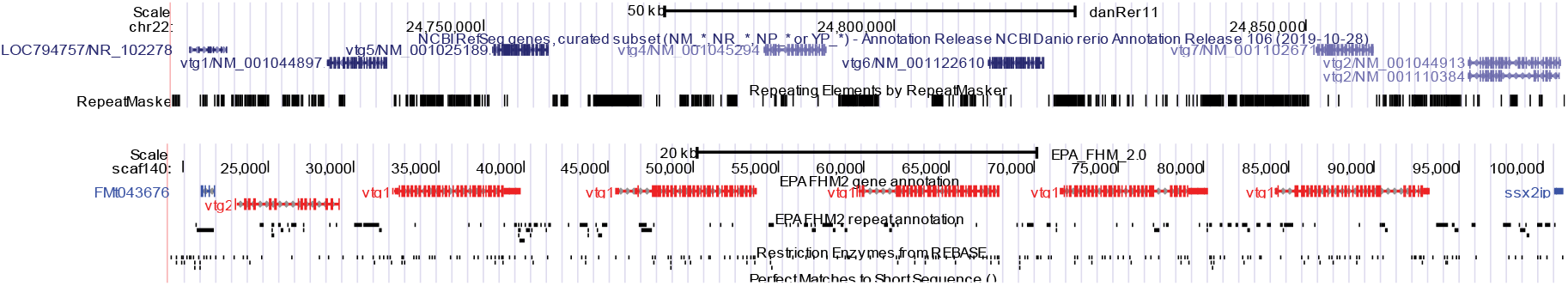
Genomic map of vitellogenin (VTG) genes in the Zebrafish and FHM. The UCSC gene map plot on the top shows the cluster of six VTG genes in the chromosome 22 of the Zebrafish genome, while the one on the bottom shows a similar cluster of six VTG genes in the scaffold 140 of the FHM2 genome.

**Figure 14:**
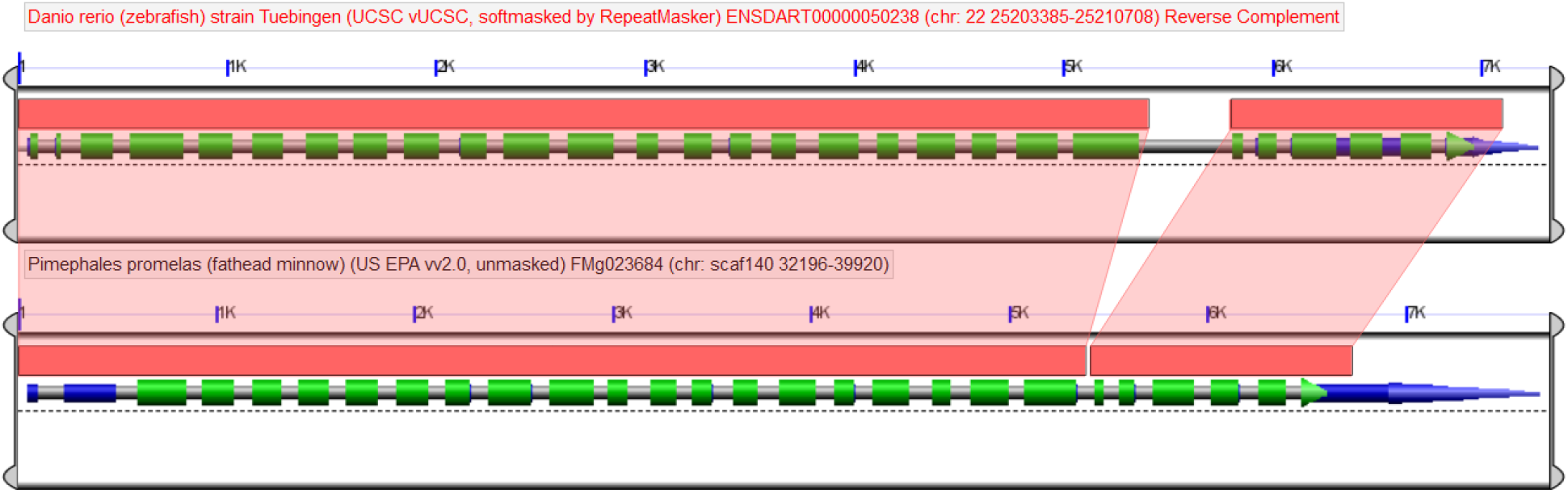
Syntenic alignment map of the VTG1 genes between zebrafish and FHM.

## DISCUSSION

We describe here the *de novo* assembly and annotation of a highly contiguous and near-complete reference genome and annotation for the FHM genome. The new reference genome, with a scaffold N50 of 12Mb and complete BUSCO score >95%, is a drastic improvement over the current FHM1 reference with respect to assembly contiguity and completeness. Several factors helped to achieve the highly contiguous FHM reference. First, we used a combination of the latest technologies to generate primary sequencing and scaffolding data for assembly. The FHM has a highly AT-rich (>60%) and repetitive genome, which is difficult, if not impossible, to be assembled with short Illumina reads alone. The FHM2 assembly relied on long PacBio reads as the primary genome assembly data, enabling assembly of long repetitive introns or intergenic regions. Illumina paired-end reads were then used to correct the relatively high sequencing error rate of PacBio reads and improve base-level accuracy of the assembly. The use of a combination of three different forms of scaffolding data, long mate-pair and fosmid Illumina reads, HiC sequencing data, and Bionano optical mapping data, enabled better connection of neighboring contigs/scaffolds and the correction of assembly/scaffolding errors during the multiple-step scaffolding process.

Second, we selected the best available software tools for assembling and scaffolding. In order to identify the best assembly tool, we evaluated several *de novo* assemblers including CANU [25, 26], FALCON/FALCON-Unzip [33], DBG2OLC [34], and Miniasm [36]. CANU was based on the well-known Celera assembler that was developed by Celera Genomics for assembling the human genome from the old Sanger sequencing data. CANU can handle both PacBio SMRT and Oxford Nanopore sequencing data. FALCON/FALCON-Unzip is a diploid assembler specifically designed for PacBio sequencing data. DBG2OLC is a hybrid assembler that takes advantage of long read PacBio data and high accuracy Illumina data. Miniasm is an OLC-based assembler for noisy long reads. Compared to the other assemblers, Miniasm is a very fast assembler but does not correct base-call errors in the assembly process. CANU and FALCON/FALCON-Unzip performed best in terms of Illumina read mapping rate, N50, and the complete BUSCOs in our test runs. In the tests, CANU achieved the highest mapping rate and the highest the complete BUSCO while FALCON/FALCON-Unzip was much faster than CANU and attained the longest contig N50, which was more than twice the N50 achieved by CANU. For these reasons, both CANU and FALCON/FALCON-Unzip were chosen as candidates to produce the final FHM genome assembly. Finally, we evaluated software parameter settings in order to optimize the overall assembly process. For example, we tested the CANU assembler with both diploid and haploid genome assembly settings, and found that CANU performed better in the diploid assembly mode than in the haploid mode based on three evaluation criteria: N50, read mapping rate and complete/single BUSCO coverage. Using the same evaluation criteria, we developed a customized scaffolding process, instead of following the established scaffolding routine (scaffolding first with mate-pair reads, followed by optical mapping data, and finally with HiC data) used by many published genome assembly studies [91–94], to accomplish the best genome assembly. Particularly, our procedure ran HiC scaffolding twice as the first and last steps in scaffolding process. The first HiC-scaffolding mainly used short-distance interaction data while the last one primarily utilized long-distance interactions in the HiC data. By using Illumina and optical mapping data as the intermediate scaffolding steps, we were able to correct scaffolding errors introduced during the initial HiC scaffolding.

Many genome assemblies were generated during the optimization process. Benchmark analyses were critical in the optimization of parameter settings for assembling and for selecting the final FHM genome assembly. As discussed above, our evaluation of genome assembly quality was based on three benchmarking analyses: N50 for assessing genome contiguity, BUSCO analysis for completeness, and Illumina read mapping rate analysis for correctness. The results from these three criteria, however, were not always consistent, making the selection of the best assembly a non-trivial problem. For example, in the comparison between one FALCON assembly and the corresponding CANU assembly, FALCON had much longer contig N50 at 628,999 than CANU of which the N50 is 360,780. However, CANU had much higher BUSCOs than FALCON as the results showed that the complete BUSCOs for the CANU and FALCON assemblies are 4373 and 4302. respectively. To deal with these contradictory results, we adopted a ranking rule where single and complete BUSCOs were given the greatest weight, followed by N50 and finally read mapping rate for assembly quality comparison and selection. In addition to these three benchmarking scores, other factors were also considered, e.g., genome duplication level measured by single and complete BUSCO score, and the closeness of the assembly size to the estimated genome size 1.11Gbps based on cytological analysis [86]. For the reason, the final FHM assembly we selected is not the one with the highest complete BUSCOs (4379/95.5%), nor it has the longest scaffold N50 at 47.6 Mbps with only 34 scaffolds (the assembly is available upon request), but is one with the highest single and complete BUSCOs with the assembly size 1.07Gpbs, which is acceptably close to the previous estimate.

With the complete BUSCOs over 95.1% and N50 at 12.0Mbps, the FHM2 reference genome was far more contiguous and complete than the FHM1 assembly. Nevertheless, the FHM2 reference genome was not fully complete and could be further improved. 13.2% of the genome is still represented by gaps though this is much smaller compared to the 33.5% gaps found in the FHM1 reference. Since most of these gap regions are very likely of highly AT-rich repetitive regions, closing these gaps require longer reads and/or deeper coverage of sequencing data from the latest third-generation sequencers, e.g., PacBio or Oxford Nanopore. Also, many scaffolds of the FHM2 reference are still at the sub-chromosome level. One likely reason is that we did not have enough tissue sample from a single fish to generate all data for genome assembly and scaffolding. So instead we used two different fish, one for generating PacBio and Illumina reads, and the other fish for the Bionano optical mapping data and HiC scaffolding data. There was large genetic variation between the two fish, which made it difficult to scaffold the assembly to the full chromosome level, however, this does have the advantage of FHM2 assembly acting as a consensus reference which benefits downstream data analysis.

We were able to comprehensively annotate the FHM2 reference genome, providing new details about its composition. In our repeat analysis, 19.3% of the FHM genome was found to be interspersed repeats by using only the known repeat libraries (data not shown) while 38.9% of the genome was identified to be interspersed repeats when combining *de novo* repeat discovery and known-library search. The result suggests that many interspersed repeats are new and may be unique to the FHM genome. In addition, our repeat analysis results show that FHM has a substantially higher proportion of simple repeats or low complexity regions compared to other teleost species, a distinguishing feature of the FHM genome.

There is great interest in using the genome assembly for the interpretation of transcriptional and epigenetic data, to this end we annotated 26,150 protein coding genes. The completeness of the protein gene annotation is indicated by the high percentage of complete BUSCOs for both the transcript (94.7%) and the genome (95.1%). The number of annotated protein coding genes in the FHM is very similar to the well-annotated zebrafish reference genome (25,638), which also has similar BUSCO scores and mapping rates. Given that the zebrafish genome is likely the most well-characterized published fish genome and is closely related to the FHM, this comparison provides strong evidence that our gene prediction and annotation is relatively complete and of high quality. Nevertheless, the UTR annotation may be incomplete as many gene models are without any UTR. We are currently working to address this issue using full-length mRNA sequencing, which will also serve to further improve gene annotation. Our annotation of small RNAs offer a glimpse of the scale of post-transcription regulations in the FHM and their potential differences among difference species. Limited by performance of prediction tools, our small RNAs annotation, especially miRNAs and piRNAs, remains preliminary at this point. Further improvement could be made with additional smRNA-seq data and experimental functional validation in future.

The comparison between the FHM and ZF reference genomes confirmed that FHM and ZF are the closest related fish species reported by the published phylogenetic study of the more than 20 vertebrates on evolution distance of neurokinin B (NKB) receptor genes (Tachykinin 3a, tac3a)[95]. This is supported by three observations from the study: 1) large syntenic genome regions shared between the two species, far more than that shared with other teleost fish species (results not shown here), 2) the majority (93%) of FHM transcripts got top BLAST-hits from the ZF genome among 27 landmark species, and 3) the high similarity of the large vitellogenin gene in gene sequence level, number of paralogs, and their chromosome arrangement. Despite the close evolutionary relationship between FHM and ZF, we found that FHM is a clearly distinguished species from ZF with respect to the following aspects: 1) large difference in GC content in intron regions with ZF at 50.6% while FHM only at 37.1%, 2) a substantial number (3350 or 7%) of FHM transcripts without any good homologous match in ZF, and 3) more compact gene structure, i.e., shorter more AT-rich intron/intergenic regions, in the FHM than ZF. All the evidence suggests that FHM and ZF have diverged substantially from their last common ancestor.

In conclusion, we *de novo* assembled a highly contiguous and complete reference genome of the fathead minnow and provided detailed annotation of the genome. Our study reveals that the FHM genome has generally much more compact gene structure than the closely related ZF though there is a high-degree similarity in coding genes between them. The new FHM reference genome, together with the comprehensive annotation, represent a major advance in environmental genome research, providing a robust framework for gene expression and regulation studies, opening the door for a much deeper understanding of changes going on at the molecular level in response to toxins and other environmental stressors.

## DATA AVAILABILITY

The FHM genome assembly and annotation data have been submitted to the NCBI Genome database with the accession number WIOS00000000. All raw DNA and RNA sequencing data associated with the project have been deposited to NCBI under the BioProject PRJNA565199, which includes 30 BioSamples (ID# SAMN12766224-SAMN12766252, and SAMN12875914), 103 SRA datasets (accesion#: SRR10135744 - SRR10135767, SRR10199005-SRR10199006, SRR10536067-SRR10536141, and SRR10613469 - SRR10613470), and one supplementary dataset (accession#SUPPF_0000003192).

## AUTHORS’ CONTRIBUTIONS

A.B. and M.K. initialized the project; A.B. supervised the project; M.K. and W.H. oversaw bioinformatics analyses. All authors were involved in planning and experimental design. D.B., M.K., R.F., M.S., and D.L. carried out the wet-lab experiments. W.H. performed genome assembly, repeat analysis, gene name and GO annotation, synteny analysis, and data visualization. J.M. developed protein-coding gene models, and J.M., M.K. and W.H. carried out gene prediction analysis and assessment; J.M. and W.H. performed comparative analyses to ZF genome/proteins/transcripts. G.T. carried out small RNAs prediction analysis. J.M., M.K., G.T. and W.H. did NCBI data submission. W.H. drafted the initial manuscript, and all authors contributed to the writing of the manuscript. All authors approved the final manuscript.

## ACKNOWLEDGMENTS

We would like to acknowledge the RTSF Genomics Core at Michigan State University for DNA and RNA sequencing services, the Dovetail Genomics for the HiC library preparation and sequencing service, and McDonnell Genome Institute at Washington University School of Medicine for the Bionano data service for the work presented in this manuscript. We would like also to thank Tom Purucker at EPA and Xuting Wang at NIEHS for EPA internal review of the manuscript.

## DISCLAIMER

The views expressed in this article are those of the authors and do not necessarily reflect the views or policies of the U.S. Environmental Protection Agency. Any mention of trade names, products, or services does not imply an endorsement by the U.S. Government or the U.S. Environmental Protection Agency (EPA). The EPA does not endorse any commercial products, services, or enterprises.

## REFERENCES

1. Ankley GT, Johnson RD: Small fish models for identifying and assessing the effects of endocrine-disrupting chemicals. ILAR J 2004, 45:469–483.

2. Ankley GT, Jensen KM, Kahl MD, Korte JJ, Makynen EA: Description and evaluation of a short-term reproduction test with the fathead minnow (Pimephales promelas). Environ Toxicol Chem 2001, 20:1276–1290.

3. Ankley GT, Kuehl DW, Kahl MD, Jensen KM, Linnum A, Leino RL, Villeneuvet DA: Reproductive and developmental toxicity and bioconcentration of perfluorooctanesulfonate in a partial life-cycle test with the fathead minnow (Pimephales promelas). Environ Toxicol Chem 2005, 24:2316–2324.

4. Geis SW, Fleming K, Mager A, Reynolds L: Modifications to the fathead minnow (Pimephales promelas) chronic test method to remove mortality due to pathogenic organisms. Environ Toxicol Chem 2003, 22:2400–2404.

5. Roush KS, Krzykwa JC, Malmquist JA, Stephens DA, Sellin Jeffries MK: Enhancing the fathead minnow fish embryo toxicity test: Optimizing embryo production and assessing the utility of additional test endpoints. Ecotoxicol Environ Saf 2018, 153:45–53.

6. Hanson C, Cairns J, Wang L, Sinha S: Principled multi-omic analysis reveals gene regulatory mechanisms of phenotype variation. Genome Res 2018, 28:1207–1216.

7. Ladd-Acosta C, Fallin MD: The role of epigenetics in genetic and environmental epidemiology. Epigenomics 2016, 8:271–283.

8. Wang RL, Biales AD, Garcia-Reyero N, Perkins EJ, Villeneuve DL, Ankley GT, Bencic DC: Fish connectivity mapping: linking chemical stressors by their mechanisms of action-driven transcriptomic profiles. BMC Genomics 2016, 17:84.

9. Brum AM, van de Peppel J, van der Leije CS, Schreuders-Koedam M, Eijken M, van der Eerden BC, van Leeuwen JP: Connectivity Map-based discovery of parbendazole reveals targetable human osteogenic pathway. Proc Natl Acad Sci U S A 2015, 112:12711–12716.

10. Exner R, Bago-Horvath Z, Bartsch R, Mittlboeck M, Retel VP, Fitzal F, Rudas M, Singer C, Pfeiler G, Gnant M, et al: The multigene signature MammaPrint impacts on multidisciplinary team decisions in ER+, HER2-early breast cancer. Br J Cancer 2014, 111:837–842.

11. Gust KA, Wilbanks MS, Guan X, Pirooznia M, Habib T, Yoo L, Wintz H, Vulpe CD, Perkins EJ: Investigations of transcript expression in fathead minnow (Pimephales promelas) brain tissue reveal toxicological impacts of RDX exposure. Aquat Toxicol 2011, 101:135–145.

12. Thomas MA, Joshi PP, Klaper RD: Gene-class analysis of expression patterns induced by psychoactive pharmaceutical exposure in fathead minnow (Pimephales promelas) indicates induction of neuronal systems. Comp Biochem Physiol C Toxicol Pharmacol 2012, 155:109–120.

13. Werner J, Ouellet JD, Cheng CS, Ju YJ, Law RD: Pulp and paper mill effluents induce distinct gene expression changes linked to androgenic and estrogenic responses in the fathead minnow (Pimephales promelas). Environ Toxicol Chem 2010, 29:430–439.

14. Biales AD, Kostich MS, Batt AL, See MJ, Flick RW, Gordon DA, Lazorchak JM, Bencic DC: Initial development of a multigene ‘omics-based exposure biomarker for pyrethroid pesticides. Aquat Toxicol 2016, 179:27–35.

15. Reuter JA, Spacek DV, Snyder MP: High-throughput sequencing technologies. Mol Cell 2015, 58:586–597.

16. Villasenor-Altamirano AB, Watson JD, Prokopec SD, Yao CQ, Boutros PC, Pohjanvirta R, Valdes-Flores J, Elizondo G: 2,3,7,8-Tetrachlorodibenzo-p-dioxin modifies alternative splicing in mouse liver. PLoS One 2019, 14:e0219747.

17. Lev Maor G, Yearim A, Ast G: The alternative role of DNA methylation in splicing regulation. Trends Genet 2015, 31:274–280.

18. Wan J, Oliver VF, Wang G, Zhu H, Zack DJ, Merbs SL, Qian J: Characterization of tissue-specific differential DNA methylation suggests distinct modes of positive and negative gene expression regulation. BMC Genomics 2015, 16:49.

19. Lu M, Zhan X: The crucial role of multiomic approach in cancer research and clinically relevant outcomes. EPMA J 2018, 9:77–102.

20. Saari TW, Schroeder AL, Ankley GT, Villeneuve DL: First-generation annotations for the fathead minnow (Pimephales promelas) genome. Environ Toxicol Chem 2017, 36:3436–3442.

21. Burns FR, Cogburn AL, Ankley GT, Villeneuve DL, Waits E, Chang YJ, Llaca V, Deschamps SD, Jackson RE, Hoke RA: Sequencing and de novo draft assemblies of a fathead minnow (Pimephales promelas) reference genome. Environ Toxicol Chem 2016, 35:212–217.

22. Florea L, Souvorov A, Kalbfleisch TS, Salzberg SL: Genome assembly has a major impact on gene content: a comparison of annotation in two Bos taurus assemblies. PLoS One 2011, 6:e21400.

23. Howe K, Clark MD, Torroja CF, Torrance J, Berthelot C, Muffato M, Collins JE, Humphray S, McLaren K, Matthews L, et al: The zebrafish reference genome sequence and its relationship to the human genome. Nature 2013, 496:498–503.

24. Lieberman-Aiden E, van Berkum NL, Williams L, Imakaev M, Ragoczy T, Telling A, Amit I, Lajoie BR, Sabo PJ, Dorschner MO, et al: Comprehensive mapping of long-range interactions reveals folding principles of the human genome. Science 2009, 326:289–293.

25. Koren S, Rhie A, Walenz BP, Dilthey AT, Bickhart DM, Kingan SB, Hiendleder S, Williams JL, Smith TPL, Phillippy AM: De novo assembly of haplotype-resolved genomes with trio binning. Nat Biotechnol 2018.

26. Koren S, Walenz BP, Berlin K, Miller JR, Bergman NH, Phillippy AM: Canu: scalable and accurate long-read assembly via adaptive k-mer weighting and repeat separation. Genome Res 2017, 27:722–736.

27. Walker BJ, Abeel T, Shea T, Priest M, Abouelliel A, Sakthikumar S, Cuomo CA, Zeng Q, Wortman J, Young SK, Earl AM: Pilon: an integrated tool for comprehensive microbial variant detection and genome assembly improvement. PLoS One 2014, 9:e112963.

28. Li H, Durbin R: Fast and accurate short read alignment with Burrows-Wheeler transform. Bioinformatics 2009, 25:1754–1760.

29. Li H, Handsaker B, Wysoker A, Fennell T, Ruan J, Homer N, Marth G, Abecasis G, Durbin R, Genome Project Data Processing S: The Sequence Alignment/Map format and SAMtools. Bioinformatics 2009, 25:2078–2079.

30. Roach MJ, Schmidt SA, Borneman AR: Purge Haplotigs: allelic contig reassignment for third-gen diploid genome assemblies. BMC Bioinformatics 2018, 19:460.

31. Ghurye J, Rhie A, Walenz BP, Schmitt A, Selvaraj S, Pop M, Phillippy AM, Koren S: Integrating Hi-C links with assembly graphs for chromosome-scale assembly. 2019:261149.

32. Boetzer M, Henkel CV, Jansen HJ, Butler D, Pirovano W: Scaffolding pre-assembled contigs using SSPACE. Bioinformatics 2011, 27:578–579.

33. Chin CS, Peluso P, Sedlazeck FJ, Nattestad M, Concepcion GT, Clum A, Dunn C, O’Malley R, Figueroa-Balderas R, Morales-Cruz A, et al: Phased diploid genome assembly with single-molecule real-time sequencing. Nat Methods 2016, 13:1050–1054.

34. Ye C, Hill CM, Wu S, Ruan J, Ma ZS: DBG2OLC: Efficient Assembly of Large Genomes Using Long Erroneous Reads of the Third Generation Sequencing Technologies. Sci Rep 2016, 6:31900.

35. Grohme MA, Schloissnig S, Rozanski A, Pippel M, Young GR, Winkler S, Brandl H, Henry I, Dahl A, Powell S, et al: The genome of Schmidtea mediterranea and the evolution of core cellular mechanisms. Nature 2018, 554:56–61.

36. Li H: Minimap and miniasm: fast mapping and de novo assembly for noisy long sequences. Bioinformatics 2016, 32:2103–2110.

37. Sahlin K, Vezzi F, Nystedt B, Lundeberg J, Arvestad L: BESST--efficient scaffolding of large fragmented assemblies. BMC Bioinformatics 2014, 15:281.

38. Luo J, Wang J, Zhang Z, Li M, Wu FX: BOSS: a novel scaffolding algorithm based on an optimized scaffold graph. Bioinformatics 2017, 33:169–176.

39. Gao S, Bertrand D, Chia BK, Nagarajan N: OPERA-LG: efficient and exact scaffolding of large, repeat-rich eukaryotic genomes with performance guarantees. Genome Biol 2016, 17:102.

40. Simao FA, Waterhouse RM, Ioannidis P, Kriventseva EV, Zdobnov EM: BUSCO: assessing genome assembly and annotation completeness with single-copy orthologs. Bioinformatics 2015, 31:3210–3212.

41. Waterhouse RM, Seppey M, Simao FA, Manni M, Ioannidis P, Klioutchnikov G, Kriventseva EV, Zdobnov EM: BUSCO applications from quality assessments to gene prediction and phylogenomics. Mol Biol Evol 2017.

42. Flynn JM, Hubley R, Goubert C, Rosen J, Clark AG, Feschotte C, Smit AF: RepeatModeler2: automated genomic discovery of transposable element families. bioRxiv 2019:856591.

43. Price AL, Jones NC, Pevzner PA: De novo identification of repeat families in large genomes. Bioinformatics 2005, 21 Suppl 1:i351–358.

44. Bao Z, Eddy SR: Automated de novo identification of repeat sequence families in sequenced genomes. Genome Res 2002, 12:1269–1276.

45. Ellinghaus D, Kurtz S, Willhoeft U: LTRharvest, an efficient and flexible software for de novo detection of LTR retrotransposons. BMC Bioinformatics 2008, 9:18.

46. Ou S, Jiang N: LTR_retriever: A Highly Accurate and Sensitive Program for Identification of Long Terminal Repeat Retrotransposons. Plant Physiol 2018, 176:1410–1422.

47. Li W, Godzik A: Cd-hit: a fast program for clustering and comparing large sets of protein or nucleotide sequences. Bioinformatics 2006, 22:1658–1659.

48. Katoh K, Misawa K, Kuma K, Miyata T: MAFFT: a novel method for rapid multiple sequence alignment based on fast Fourier transform. Nucleic Acids Res 2002, 30:3059–3066.

49. Wheeler TJ: Large-Scale Neighbor-Joining with NINJA. In; Berlin, Heidelberg. Springer Berlin Heidelberg; 2009: 375–389.

50. Benson G: Tandem repeats finder: a program to analyze DNA sequences. Nucleic Acids Res 1999, 27:573–580.

51. Hubley R, Finn RD, Clements J, Eddy SR, Jones TA, Bao W, Smit AF, Wheeler TJ: The Dfam database of repetitive DNA families. Nucleic Acids Res 2016, 44:D81–89.

52. Bao W, Kojima KK, Kohany O: Repbase Update, a database of repetitive elements in eukaryotic genomes. Mob DNA 2015, 6:11.

53. Stanke M, Diekhans M, Baertsch R, Haussler D: Using native and syntenically mapped cDNA alignments to improve de novo gene finding. Bioinformatics 2008, 24:637–644.

54. Grabherr MG, Haas BJ, Yassour M, Levin JZ, Thompson DA, Amit I, Adiconis X, Fan L, Raychowdhury R, Zeng Q, et al: Full-length transcriptome assembly from RNA-Seq data without a reference genome. Nat Biotechnol 2011, 29:644–652.

55. Song L, Florea L: Rcorrector: efficient and accurate error correction for Illumina RNA-seq reads. Gigascience 2015, 4:48.

56. [github.com/FelixKrueger/TrimGalore]

57. Martin M: Cutadapt removes adapter sequences from high-throughput-seqeuncing reads. EMBnetjournal 2011, 17:10–12.

58. Dobin A, Davis CA, Schlesinger F, Drenkow J, Zaleski C, Jha S, Batut P, Chaisson M, Gingeras TR: STAR: ultrafast universal RNA-seq aligner. Bioinformatics 2013, 29:15–21.

59. Cantarel BL, Korf I, Robb SMC, Parra G, Ross E, Moore B, Holt C, Sanchez Alvarado A, Yandell M: MAKER: An easy-to-use annotation pipeline designed for emerging model organism genomes. Genome Res 2008, 18.

60. Haas BJ, Salzberg SL, Zhu W, Pertea M, Allen JE, Orvis J, White O, Buell CR, Wortman JR: Automated eukaryotic gene structure annotation using EVidenceModeler and the Program to Assemble Spliced Alignments. Genome Biol 2008, 9:R7.

61. Stanke M, Keller O, Gunduz I, Hayes A, Waack S, Morgenstern B: AUGUSTUS: ab initio prediction of alternative transcripts. Nucleic Acids Res 2006, 34:W435–439.

62. Korf I: Gene finding in novel genomes. BMC Bioinformatics 2004, 5.

63. Lukashin AV, Borodovsky M: GeneMark.hmm: new solutions for gene finding. 1998, 26:1107–1115.

64. Kent WJ: BLAT--the BLAST-like alignment tool. Genome Res 2002, 12:656–664.

65. Wu TD, Watanabe CK: GMAP: a genomic mapping and alignment program for mRNA and EST sequences. Bioinformatics 2005, 21:1859–1875.

66. Haas BJ, Papanicolaou A, Yassour M, Grabherr M, Blood PD, Bowden J, Couger MB, Eccles D, Li B, Lieber M, et al: De novo transcript sequence reconstruction from RNA-seq using the Trinity platform for reference generation and analysis. Nature protocols 2013, 8:1494–1512.

67. Edgar RC: Search and clustering orders of magnitude faster than BLAST. Bioinformatics 2010, 26:2460–2461.

68. Bushnell B: BBMap. 2014.

69. Langmead B, Salzberg SL: Fast gapped-read alignment with Bowtie 2. Nat Methods 2012, 9.

70. Buchfink B, Xie C, Huson DH: Fast and sensitive protein alignment using DIAMOND. Nat Methods 2015, 12:59–60.

71. Altschul SF, Gish W, Miller W, Meyers EW, Lipman DJ: Basic Local Alignment Search Tool. Journal of Molecular Biology 1990, 215.

72. Camacho C, Coulouris G, Avagyan V, Ma N, Papadopoulos J, Bealer K, Madden TL: BLAST+: architecture and applications. BMC Bioinformatics 2009, 10:421.

73. Pertea G, Pertea M: GFF Utilities: GffRead and GffCompare. F1000Res 2020, 9.

74. Chan PP, Lowe TM: tRNAscan-SE: Searching for tRNA Genes in Genomic Sequences. Methods Mol Biol 2019, 1962:1–14.

75. An J, Lai J, Lehman ML, Nelson CC: miRDeep*: an integrated application tool for miRNA identification from RNA sequencing data. Nucleic Acids Res 2013, 41:727–737.

76. Wang K, Liang C, Liu J, Xiao H, Huang S, Xu J, Li F: Prediction of piRNAs using transposon interaction and a support vector machine. BMC Bioinformatics 2014, 15:419.

77. Wang K, Hoeksema J, Liang C: piRNN: deep learning algorithm for piRNA prediction. PeerJ 2018, 6:e5429.

78. Kalvari I, Nawrocki EP, Argasinska J, Quinones-Olvera N, Finn RD, Bateman A, Petrov AI: Non-Coding RNA Analysis Using the Rfam Database. Curr Protoc Bioinformatics 2018, 62:e51.

79. Jones P, Binns D, Chang HY, Fraser M, Li W, McAnulla C, McWilliam H, Maslen J, Mitchell A, Nuka G, et al: InterProScan 5: genome-scale protein function classification. Bioinformatics 2014, 30:1236–1240.

80. Gotz S, Garcia-Gomez JM, Terol J, Williams TD, Nagaraj SH, Nueda MJ, Robles M, Talon M, Dopazo J, Conesa A: High-throughput functional annotation and data mining with the Blast2GO suite. Nucleic Acids Res 2008, 36:3420–3435.

81. Lyons E, Freeling M: How to usefully compare homologous plant genes and chromosomes as DNA sequences. Plant J 2008, 53:661–673.

82. Kielbasa SM, Wan R, Sato K, Horton P, Frith MC: Adaptive seeds tame genomic sequence comparison. Genome Res 2011, 21:487–493.

83. Tang H, Lyons E, Pedersen B, Schnable JC, Paterson AH, Freeling M: Screening synteny blocks in pairwise genome comparisons through integer programming. BMC Bioinformatics 2011, 12:102.

84. Lyons E, Freeling M, Kustu S, Inwood W: Using genomic sequencing for classical genetics in E. coli K12. PLoS One 2011, 6:e16717.

85. Haug-Baltzell A, Stephens SA, Davey S, Scheidegger CE, Lyons E: SynMap2 and SynMap3D: web-based whole-genome synteny browsers. Bioinformatics 2017, 33:2197–2198.

86. Gold JR, Amemiya CT: Genome size variation in North American minnows (Cyprinidae). II. Variation among 20 species. Genome 1987, 29:481–489.

87. Gao B, Shen D, Xue S, Chen C, Cui H, Song C: The contribution of transposable elements to size variations between four teleost genomes. Mob DNA 2016, 7:4.

88. Altschul SF, Gish W, Miller W, Myers EW, Lipman DJ: Basic local alignment search tool. J Mol Biol 1990, 215:403–410.

89. Ozata DM, Gainetdinov I, Zoch A, O’Carroll D, Zamore PD: PIWI-interacting RNAs: small RNAs with big functions. Nat Rev Genet 2019, 20:89–108.

90. Nelson ADL, Haug-Baltzell AK, Davey S, Gregory BD, Lyons E: EPIC-CoGe: managing and analyzing genomic data. Bioinformatics 2018, 34:2651–2653.

91. Du H, Yu Y, Ma Y, Gao Q, Cao Y, Chen Z, Ma B, Qi M, Li Y, Zhao X, et al: Sequencing and de novo assembly of a near complete indica rice genome. Nat Commun 2017, 8:15324.

92. Zhang L, Li S, Luo J, Du P, Wu L, Li Y, Zhu X, Wang L, Zhang S, Cui J: Chromosome-level genome assembly of the predator Propylea japonica to understand its tolerance to insecticides and high temperatures. Mol Ecol Resour 2020, 20:292–307.

93. Field MA, Rosen BD, Dudchenko O, Chan EKF, Minoche AE, Edwards RJ, Barton K, Lyons RJ, Tuipulotu DE, Hayes VM, et al: Canfam_GSD: De novo chromosome-length genome assembly of the German Shepherd Dog (Canis lupus familiaris) using a combination of long reads, optical mapping, and Hi-C. Gigascience 2020, 9.

94. Shi J, Ma X, Zhang J, Zhou Y, Liu M, Huang L, Sun S, Zhang X, Gao X, Zhan W, et al: Chromosome conformation capture resolved near complete genome assembly of broomcorn millet. Nat Commun 2019, 10:464.

95. Biran J, Golan M, Mizrahi N, Ogawa S, Parhar IS, Levavi-Sivan B: Direct regulation of gonadotropin release by neurokinin B in tilapia (Oreochromis niloticus). Endocrinology 2014, 155:4831–4842.

