## Supplemental Materials for "De novo assembly and annotation of a highly contiguous reference genome of the fathead minnow (Pimephales promelas) reveals an AT-rich repetitive genome with compact gene structure"

[3]Present Address: the Jackson laboratory for genomic medicine, Farmington, CT 06032, USA

### SUPPLEMENTAL MATERIALS

#### Supplemental Tables

*Suppl Table 1: CANU assembler parameter settings for FHM assembling with PacBio sequencing reads*

| CANU parameter | value |
|---|---:|
| **genomeSize** | 1200000000 |
| **corOutCoverage** | 200 |
| **corMaxEvidenceErate** | 0.15 |
| **corOvlErrorRate** | 0.24 |
| **obtOvlErrorRate** | 0.045 |
| **utgOvlErrorRate** | 0.045 |
| **corErrorRate** | 0.3 |
| **obtErrorRate** | 0.045 |
| **utgErrorRate** | 0.045 |
| **cnsErrorRate** | 0.075 |

Suppl Table 2: List of *software tools employed in some manner in MAKER and PASA/EVM annotation pipelines*

| Tool (version) | MAKER | PASA |
|---|---|---|
| RepeatMasker (open-4.0.7) | X | X* |
| ncbi-blast+ (2.6.0 and 2.2.31) | X | X* |
| Exonerate (2.2.2) | X | X* |
| BLAT (36x1) | | X |
| gmap (2017-11-15) | | X |
| Augustus (3.2.2) | X | X* |
| Genemark-ES (4.38) | X | X* |
| SNAP (2006-07-28) | X | X* |
| TransDecoder (5.5.0) | | X |
| Hmmer (3.1b2, vs. Pfam version 32.0;) | | X |
| EvidenceModeler (1.1.1) | | X |

*Indirect use as (primarily) outputs from these tools generated when run under MAKER were employed as inputs to “PASA” pipeline at EVidenceModeler step.

*Suppl Table 3: List of NCBI SRA sources of zebrafish RNA-seq reads used for mapping rate analysis*

| **Accession#** | **Tissue** | **#reads** | **bases(gb)** | **Read length** |
|---|---|---|---|---|
| **SRR5378555** | juvenile male testis | 23,581,077 | 4.7 | 101 |
| **SRR5022640** | embryo 10 hpf | 21,629,414 | 4 | 100 |
| **SRR1609753** | Muscle | 18,159,812 | 3.6 | 100 |
| **SRR5893042** | pooled male and female embryo | 22,375,051 | 3.4 | 76 |
| **SRR5893166** | pooled male and female embryo | 20,266,392 | 3.1 | 76 |
| **SRR891504** | Liver | 28,442,972 | 2.9 | 50 |
| **SRR6308293** | Blood | 20,461,146 | 3.8 | 98/89 |
| **SRR5893071** | pooled male and female embryo | 18,148,267 | 2.8 | 76 |
| **SRR891495** | Heart | 27,037,328 | 2.8 | 51 |
| **SRR630469** | embryo 5 dpf | 20,109,108 | 2.1 | 51 |
| **SRR5893069** | pooled male and female embryo | 15,220,175 | 2.3 | 76 |
| **SRR5342780** | Ovary | 27,797,691 | 3.9 | 70 |
| **SRR6652897** | Brain | 14,200,064 | 2.9 | 101 |
| **SRR4375303** | pooled male and female embryo | 11,840,646 | 1.8 | 76 |
| **SRR5599699** | Mixed Tissues (muscle, ovary, kidney, gill, liver, intestines, heart, brain) | 12,699,919 | 2.5 | 100 |
| **SRR891511** | Brain | 15,662,615 | 1.6 | 51 |
| **SRR4375302** | pooled male and female embryo | 8,641,032 | 1.3 | 76 |
| | Total # of read pairs | 326,272,709 | | |

Suppl Table 4: Comparison repeat element distributions of the FHM genome with the other four teleost genomes. Note that repeats statistics marked be "*" were from the published study by Gao et al.[87].

| | Fathead Minnow | | Zebrafish* | | Medaka* | | Stickleback* | | Tetraodon* | |
|---|---|---|---|---|---|---|---|---|---|---|
| | **Count** | **(%/Mb)** | **Count** | **(%/Mb)** | **Count** | **(%/Mb)** | **Count** | **(%/Mb)** | **Count** | **(%/Mb)** |
| **Total retrotransposons** | 249678 | 9.08/96.79 | 533112 | 12.00/164.29 | 215153 | 8.37/58.71 | 105276 | 6.61/29.50 | 35607 | 4.00/12.08 |
| **SINE** | 22354 | 0.25/2.71 | 136879 | 2.24/30.64 | 30578 | 0.68/4.79 | 11523 | 0.67/2.97 | 1498 | 0.09/0.26 |
| **LINE** | 78893 | 2.20/23.45 | 132888 | 3.85/52.78 | 112487 | 4.97/34.86 | 35604 | 2.60/11.61 | 19385 | 1.97/5.94 |
| **LTR** | 148431 | 6.62/70.63 | 160149 | 5.90/80.87 | 72088 | 2.72/19.05 | 58159 | 3.34/14.92 | 14724 | 1.95/5.89 |
| **DNA transposons** | 1138712 | 21.47/228.91 | 2368307 | 41.07/562.49 | 282359 | 11.00/77.14 | 73571 | 4.47/19.96 | 21901 | 1.55/4.68 |
| **Unclassified** | 583472 | 8.36/89.13 | 228249 | 3.43/46.92 | 397468 | 14.32/100.42 | 72717 | 3.14/14.02 | 18465 | 1.58/4.78 |
| **Total interspersed repeats** | | **38.90/414.83** | | **56.49/773.70** | | **33.70/236.28** | | **14.21/63.48** | | **7.13/21.55** |
| **Small RNA** | 3015 | 0.05/0.51 | 13817 | 0.12/1.65 | 7223 | 0.16/1.09 | 2950 | 0.10/0.45 | 784 | 0.04/0.13 |
| **Satellite** | 30422 | 0.51/5.49 | 75515 | 1.50/20.61 | 3046 | 0.16/1.13 | 1309 | 0.09/0.41 | 560 | 0.08/0.23 |
| **Simple repeats** | **354572** | **1.56/16.61** | **42321** | **0.99/13.50** | **15283** | **0.29/2.03** | **8876** | **0.25/1.12** | **22873** | **0.74/2.25** |
| **Low complexity** | **35283** | **0.18/1.92** | **1128** | **0.03/0.35** | **149** | **0.00/0.03** | **243** | **0.01/0.04** | **300** | **0.02/0.05** |

*Suppl Table 5: Mapping rates of 6.43M randomly selected trimmed paired-end FHM RNA-seq reads to exemplar transcripts from different model sets along the path to the final gene models*

| **Target** | **# of transcripts** | **Unique mapping rate** | **Multi mapping rate** | **Total mapping rate** |
|---|---|---|---|---|
| **Maker** | 30,909 | 71.7% | 6.2% | 77.9% |
| **PASA/EVM** | 37,190 | 72.9% | 5.1% | 77.9% |
| **Pre-filtered models** | 36,881 | 78.6% | 6.8% | 85.2% |
| **Filtered models** | 26,150 | 78.3% | 6.6% | 84.9% |
| **Final curated models** | 26,150 | 78.3% | 6.6% | 84.9% |

*Suppl Table 6: - FHM tRNA prediction statistics by tRNAscan-SE (2.0.5) in comparison with other species*

| *tRNA Summary statistics* | *FHM* | *D. rerio* | *hg38* | *G. morhua* | *G. aculeatus* | *O. niloticus* | *O. latipes* | *T. rubripes* | *T. nigroviridis* |
|---|---|---|---|---|---|---|---|---|---|
| | | | Human | Atl. Cod | Stickleback | Nile tilapia | Medaka | Fugu | Tetraodon |
| *# predicted:* | 3776 | 20581 | 853 | 2540 | 2988 | 1193 | 3056 | 691 | 659 |
| *# pseudogenes:* | 383 | 4277 | 89 | 689 | 228 | 409 | 1864 | 98 | 114 |
| *# failed 2nd cutoff:* | 480 | 4982 | 98 | 572 | 375 | 147 | 170 | 62 | 106 |
| *# failed tertiary filter:* | 209 | 2439 | | 54 | 309 | 2 | | | |
| *# after filtering:* | 2704 | 8879 | 432 | 1212 | 2067 | 633 | 1021 | 526 | 432 |
| *# mismatched isotype:* | 26 | 149 | 2 | 31 | 13 | 7 | 15 | 2 | 5 |
| *# unknown isotype:* | 3 | 5 | | 7 | 7 | | | | 2 |
| *# unexpected anticodon:* | 5 | 49 | 1 | 6 | 1 | 5 | 4 | 1 | |
| *# in high confidence set:* | 2670 | 8676 | 429 | 1168 | 2046 | 621 | 1002 | 523 | 425 |
| *Selenocysteine tRNAs (TCA)* | 1 | 3 | 1 | 1 | 1 | 1 | 1 | 1 | 1 |

*Suppl Table 7: List of FHM tRNAs by anticodon counts*

| **AA (#tRNAs)** | **tRNA codon: #tRNAs** | | | | | |
|---|---|---|---|---|---|---|
| **Ala (91)** | AGC:35 | GGC:0 | CGC:17 | TGC:39 | | |
| **Gly (204)** | ACC:0 | GCC:86 | CCC:17 | TCC:101 | | |
| **Pro (133)** | AGG:75 | GGG:0 | CGG:20 | TGG:38 | | |
| **Thr (79)** | AGT:30 | GGT:0 | CGT:9 | TGT:40 | | |
| **Val (224)** | AAC:69 | GAC:0 | CAC:84 | TAC:71 | | |
| **Ser (229)** | AGA:59 | GGA:0 | CGA:19 | TGA:46 | ACT:0 | GCT:105 |
| **Arg (102)** | ACG:23 | GCG:0 | CCG:8 | TCG:7 | CCT:38 | TCT:26 |
| **Leu (361)** | AAG:60 | GAG:0 | CAG:32 | TAG:43 | CAA:158 | TAA:68 |
| **Phe (92)** | AAA:0 | GAA:92 | | | | |
| **Asn (49)** | ATT:0 | GTT:49 | | | | |
| **Lys (104)** | CTT:47 | TTT:57 | | | | |
| **Asp (157)** | ATC:0 | GTC:157 | | | | |
| **Glu (42)** | CTC:22 | TTC:20 | | | | |
| **His (48)** | ATG:0 | GTG:48 | | | | |
| **Gln (89)** | CTG:45 | TTG:44 | | | | |
| **Ile (137)** | AAT:79 | GAT:0 | TAT:58 | | | |
| **Met (164)** | CAT:164 | | | | | |
| **Tyr (14)** | ATA:0 | GTA:14 | | | | |
| **Supres (0)** | CTA:0 | TTA:0 | | | | |
| **Cys (164)** | ACA:0 | GCA:164 | | | | |
| **Trp (186)** | CCA:186 | | | | | |
| **SelCys (1)** | TCA:1 | | | | | |

*Suppl Table 8: Comparison of #of miRNAs between FHM and other species. The number of miRNAs reported here for the other species were from mirbase.org.*

| *Species* | *# miRNAs* |
|---|---|
| *P. promelas (FHM)* | 620 |
| *D. rerio* | 374 |
| *G. morhua* | 516 |
| *O. latipes* | 146 |
| *T. rubripes* | 108 |
| *T. nigroviridis* | 109 |
| *O. niloticus* | 695 |
| *H. sapiens* | 2656 |
| *M. musculus* | 1978 |

*Suppl Table 9: List of other ncRNAs detected with Infernal/Rfam analyses*

•

| ncRNA | Total count | # families |
|---|---|---|
| **LSU_rRNA_eu** | 16 | |
| **SSU_rRNA_eu** | 47 | |
| **5S_rRNA** | 3459 | |
| **U (snRNA)** | 347 | 20 |
| **SNORNA** | 201 | 96 |
| **SCARNA** | 13 | 6 |
| **RNase_MRP** | 1 | |
| **RNaseP_nuc** | 6 | |
| **Metazo_SRP** | 43 | |

**Supplemental Figures**

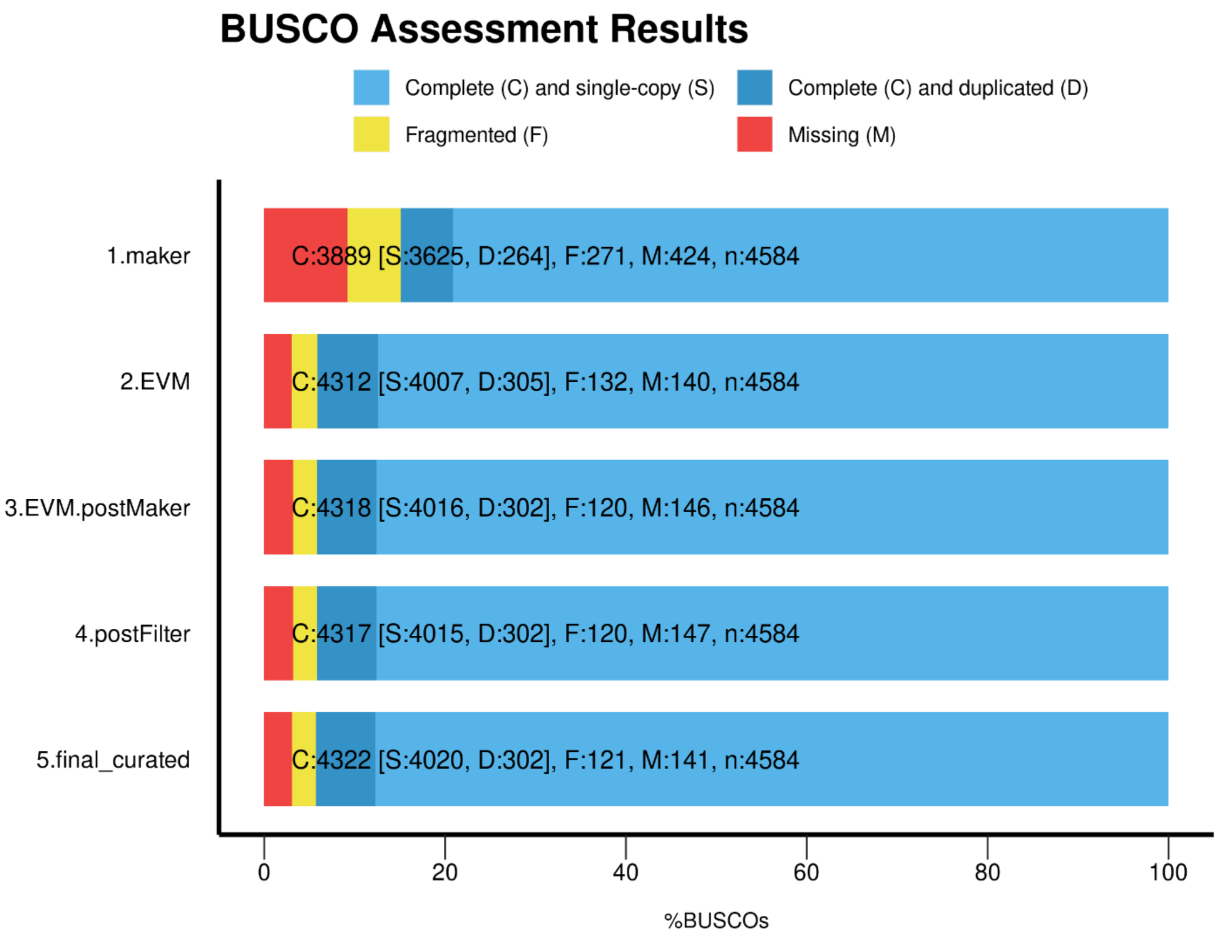


*Suppl Figure 1: Comparison of BUSCO results of different exemplar proteins sets from different steps of our gene prediction pipeline. Top two lines show results for initial outputs from Maker and PASA/EVM annotation pipelines. The third line shows the BUSCO content after having reprocessed the EVM models back through Maker. The fourth line shows the results after filtering 36,881 post Maker EVM gene models to 26,150 post-filtering models, and the fifth line shows the results after manual curation of the filtered models (i.e., the final genes/proteins).*

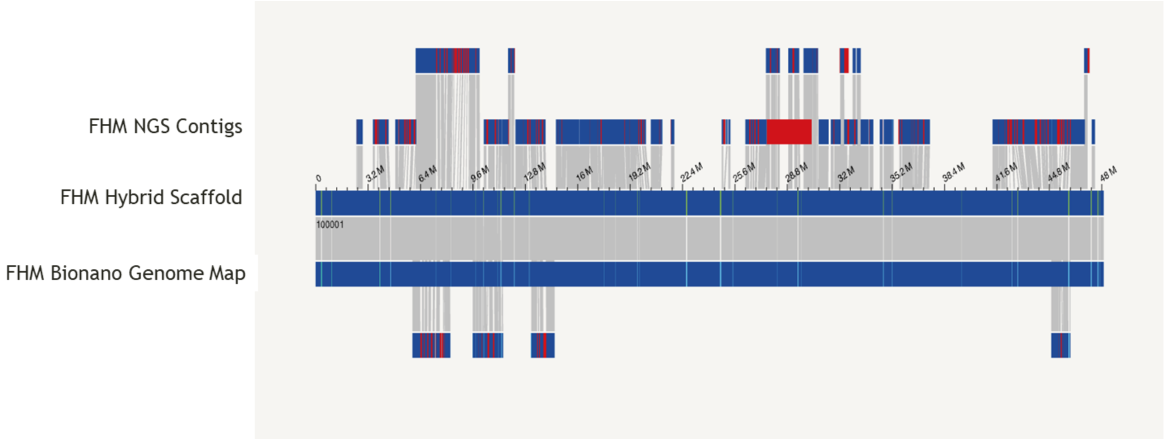


*Suppl Figure 2: Illustration of generating FHM hybrid assembly from NGS-based assembly and Bionano genome maps. The figure shows an example of generating a hybrid scaffold from NGS assembly contigs and Bionano genome maps.*

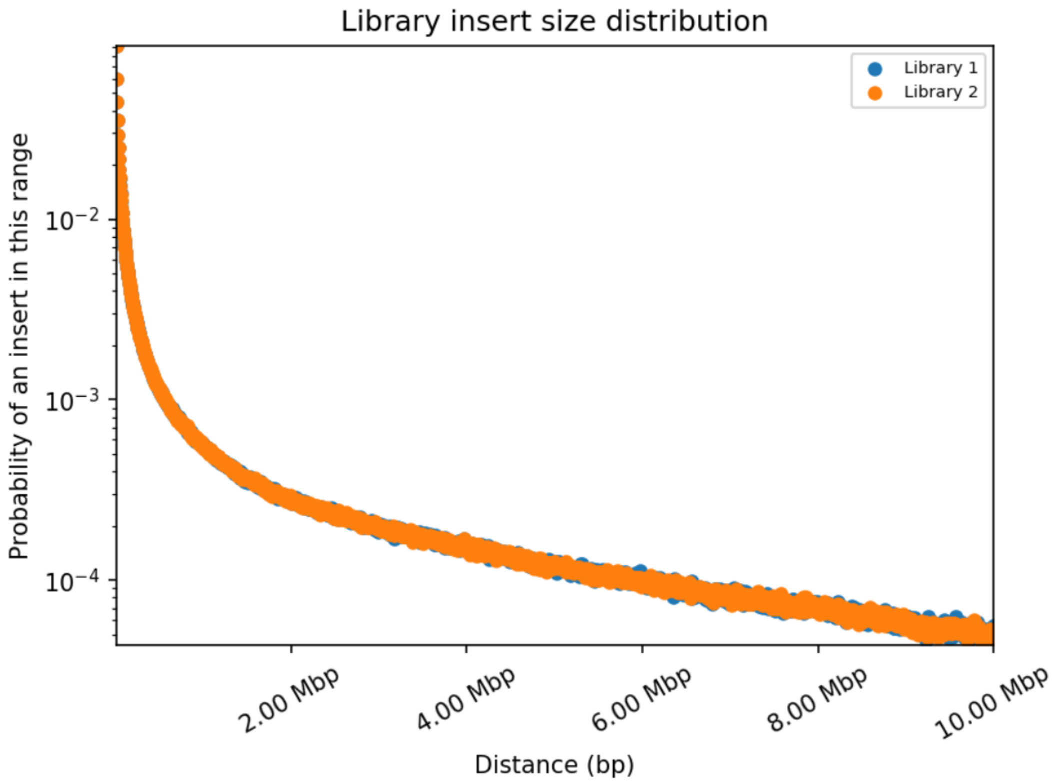


*Suppl Figure 3: library size distribution of HiC Illumina sequencing data*

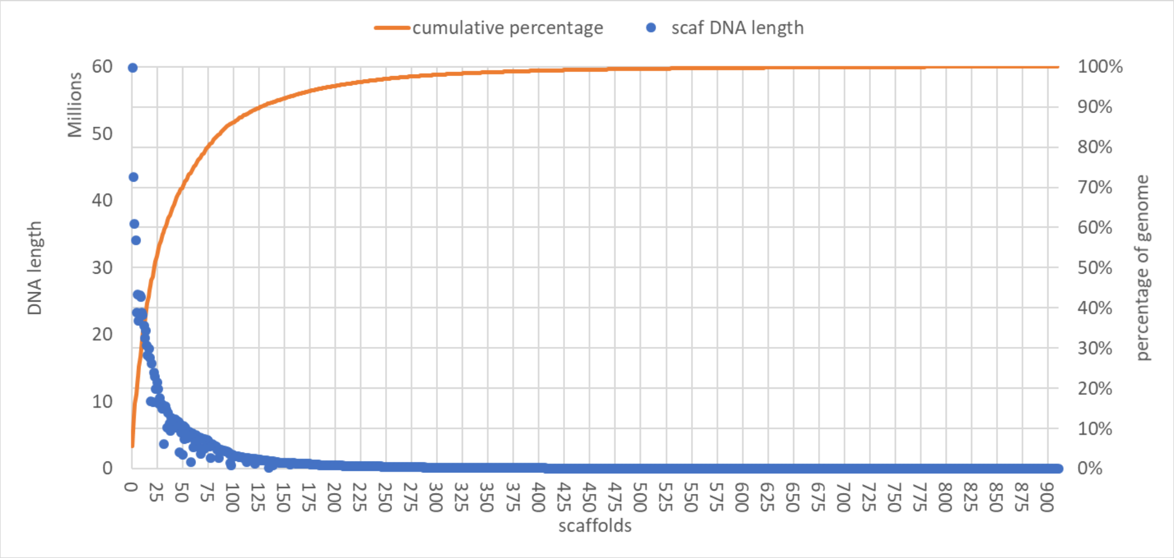


*Suppl Figure 4: Scaffold length distribution of the FHM genome assembly. The blue dots are lengths of individual scaffolds, and yellow ones are the cumulative percentage of the total genome length.*

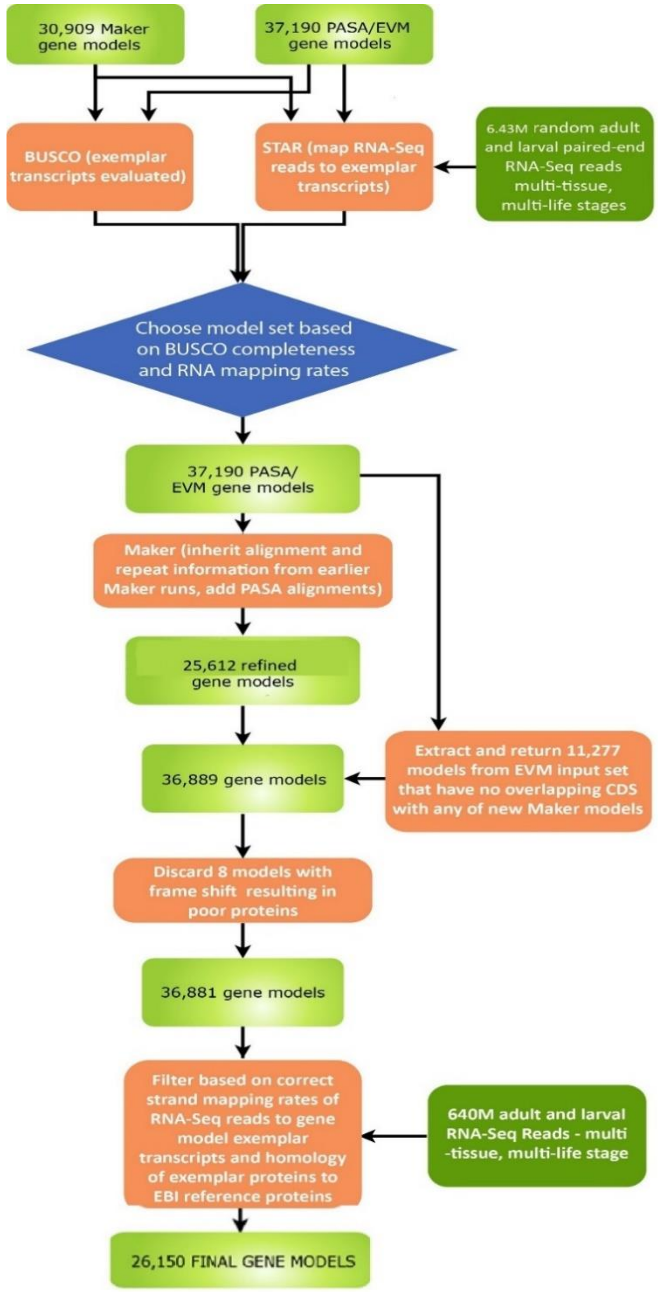


*Suppl Figure 5: Flowchart of FHM2 protein coding gene prediction following initial Maker and PASA/EVM runs*

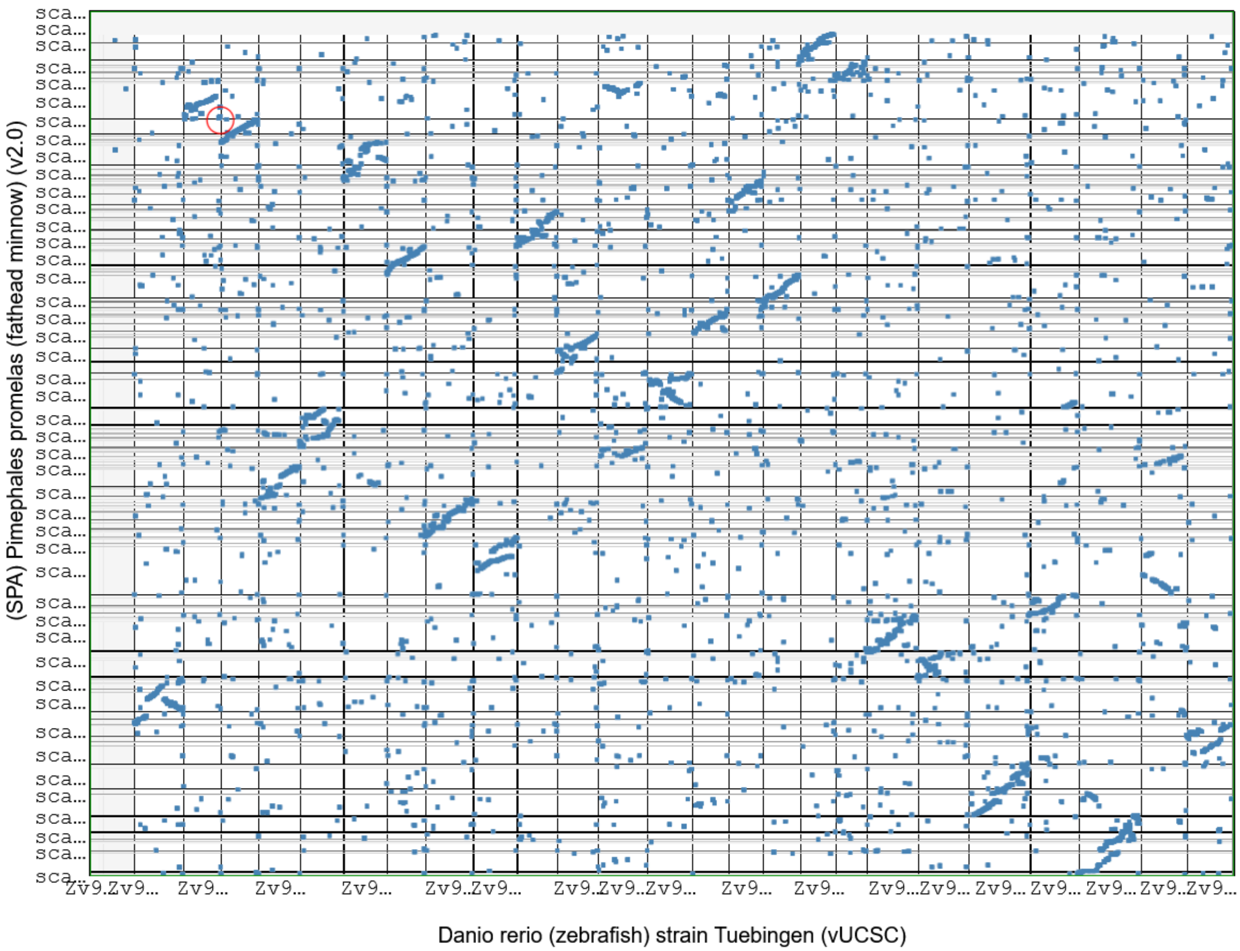

*Suppl Figure 6: Syntenic dot-plot of genomic regions of FHM and Zebrafish. X-axis is the Zebrafish reference genome and y-axis is FHM's Syntenic Path Assembly (SPA) based on synteny with zebrafish.*

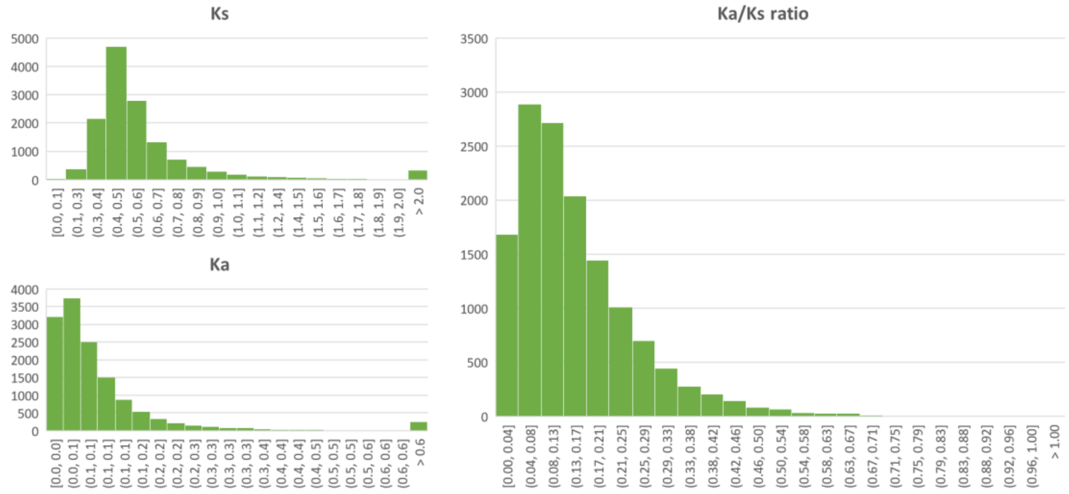


*Suppl Figure 7: Histograms of Ks, and Ka, and Ka/Ks ratio from the syntenic analysis of CDS of FHM and Zebrafish. On average, Ks is 0.62, and Ka is 0.10, and Ka/Ks ratio is 0.16, which indicates negative selection or purifying selection in the evolutionary process.*

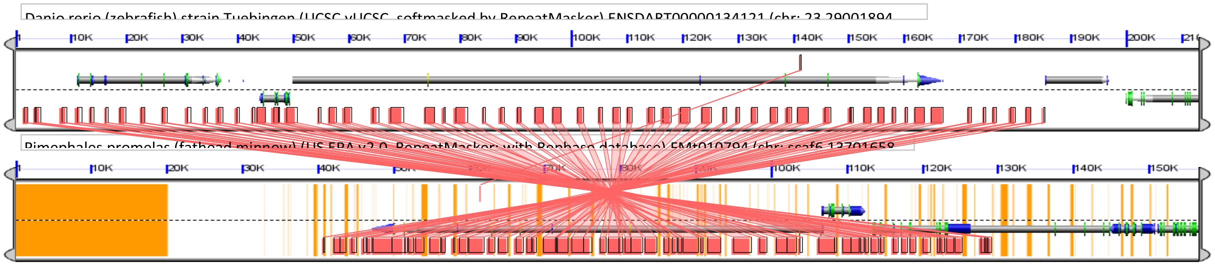


*Suppl Figure 8: Example of a syntenic region between the Zebrafish genome and FHM genome. The syntenic region in the zebrafish genome is centered around the gene ENSDART00000134121 and that of the FHM is centered around the gene FMt010794. The yellow regions in the FHM genome are of repeat-masked or gap regions. In the Zebrafish genome the syntenic region is more than 185Kb in length while that of the FHM genome is less than 90kb. For this syntenic region, the FHM genome lost over half of sequences, most of intron regions or intergenic regions, in the evolution process.*

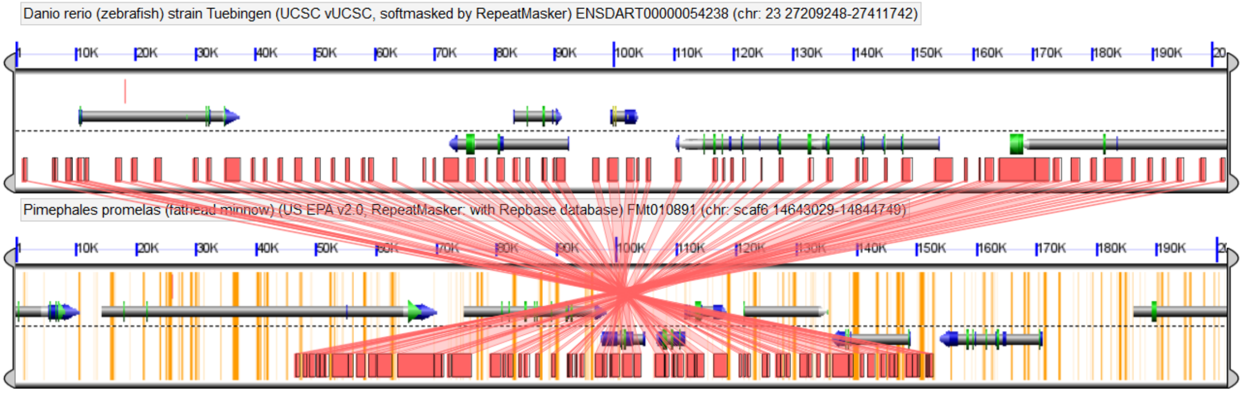


*Suppl Figure 9: GEvo syntenic map of the genomic region surrounding the FMt010891 gene in FHM and the ENDARTT00000054238491 gene in zebrafish. The genomic region containing the syntenic region in the Zebrafish is of about 200Kbps while the corresponding region in the FHM genome has only about 105Kbps, which is almost twice shorter.*

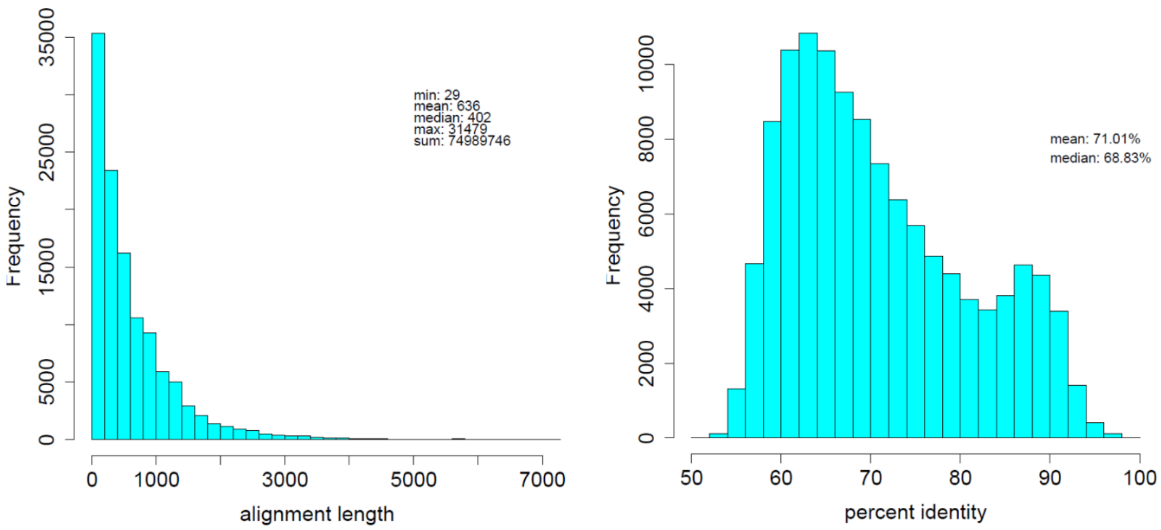


*Suppl Figure 10: Distribution of CDS HSPs between Zebrafish and FHM. The HSPs were generated by LAST in the syntenic region analysis. On the left is the histogram of alignment length and on the right is the one for percent identify of HSPs.*

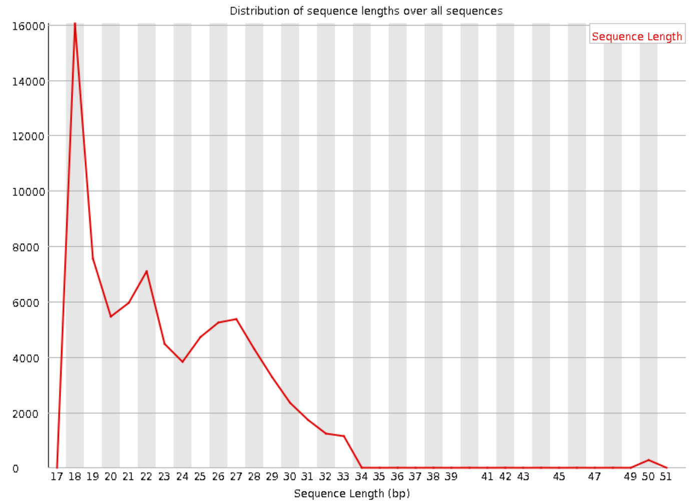


*Suppl Figure 11: Size distribution of Piano-predicted FHM piRNAs*

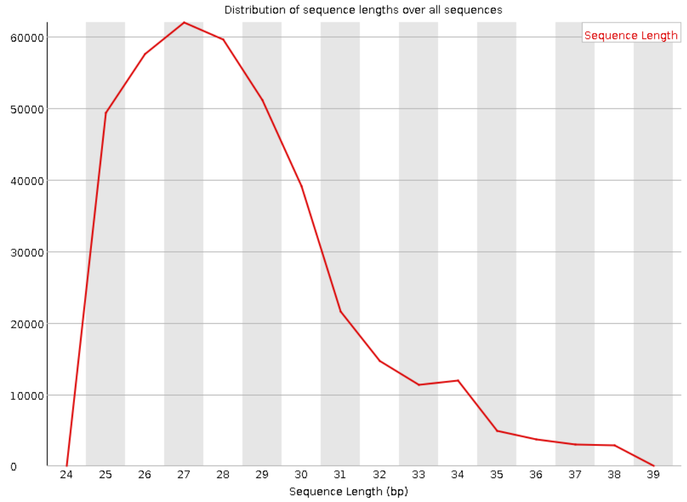


*Suppl Figure 12:  Size distribution of piRNN-predicted FHM piRNAs*
